# Learning the Rules of Cell Competition without Prior Scientific Knowledge

**DOI:** 10.1101/2021.11.24.469554

**Authors:** Christopher J. Soelistyo, Giulia Vallardi, Guillaume Charras, Alan R. Lowe

## Abstract

Deep learning is now a powerful tool in microscopy data analysis, and is routinely used for image processing applications such as segmentation and denoising. However, it has rarely been used to directly learn mechanistic models of a biological system, owing to the complexity of the internal representations. Here, we develop an end-to-end machine learning model capable of learning the rules of a complex biological phenomenon, cell competition, directly from a large corpus of time-lapse microscopy data. Cell competition is a quality control mechanism that eliminates unfit cells from a tissue and during which cell fate is thought to be determined by the local cellular neighborhood over time. To investigate this, we developed a new approach (*τ*-VAE) by coupling a probabilistic encoder to a temporal convolution network to predict the fate of each cell in an epithelium. Using the *τ*-VAE’s latent representation of the local tissue organization and the flow of information in the network, we decode the physical parameters responsible for correct prediction of fate in cell competition. Remarkably, the model autonomously learns that cell density is the single most important factor in predicting cell fate – a conclusion that is in agreement with our current understanding from over a decade of scientific research. Finally, to test the learned internal representation, we challenge the network with experiments performed in the presence of drugs that block signalling pathways involved in competition. We present a novel discriminator network that, using the predictions of the *τ*-VAE, can identify conditions which deviate from the normal behaviour, paving the way for automated, mechanism-aware drug screening.

## Introduction

Cell competition is a phenomenon that results in the elimination of less fit cells from a tissue – a critical process in development, homeostasis and disease [1]. The viability of loser cells depends strongly on context: when they are cultured alone, they thrive, but when in a mixed population, they are eliminated by cells with greater fitness (**Fig 1a**). In development, competition acts as a quality control mechanism and also participates in pattern formation [2]. In cancer, competition has been hypothesised to underlie the heterogeneity in cell types present in tumours and promote the emergence of the most aggressive cells [3]. However, the rules that determine individual cell fate are poorly understood. A number of mechanisms of cell competition have been identified to date involving either biochemical competition (for example through competition for pro-survival growth factors) or mechanical competition (for example a fast growing clone compresses cells in a slow growing clone, which results in cell extrusion for the now denser slow growing clone) [1, 4, 5]. While competition was initially thought to take place only at the interface between cell lineages, the discovery of mechanical competition revealed that this is not necessarily the case and that extrusion may take place several cell diameters away from this interface [4]. Over a decade of experimental research has suggested that local cell density is a key determinant of cell fate in mechanical competition [6, 4, 7, 8, 9]. Here, we define the term “local cell density” as the inverse of the apical surface area occupied by the cell of interest and its direct neighbours [8, 9]. However, the vast majority of studies have examined mechanisms of competition at the population level, owing to the difficulty of quantitatively describing the time evolution of the environment of each cell within a population. As such, our understanding of the cell-scale topological and physical parameters that determine fate in competition remains incomplete.

**Figure 1:**
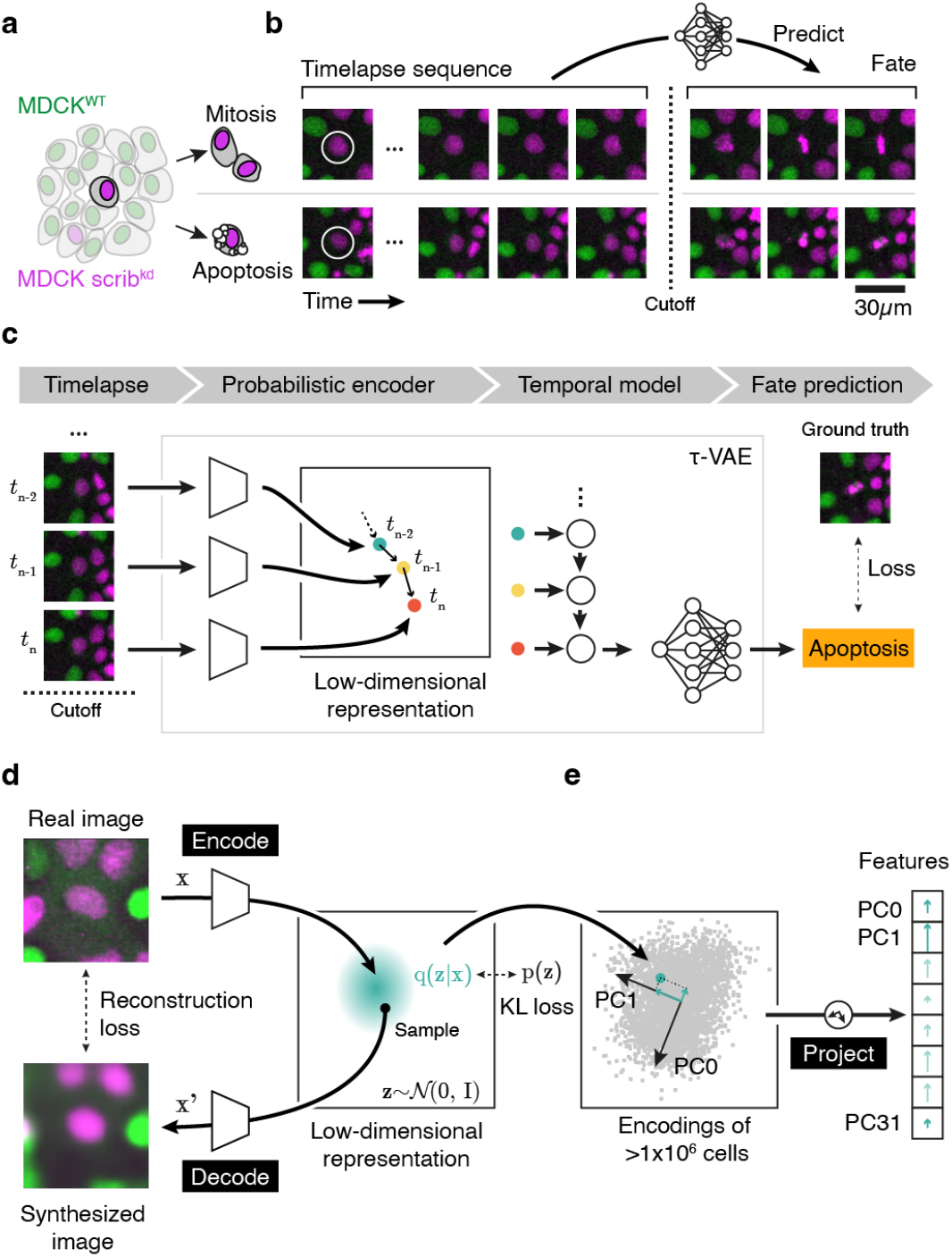
Learning a meaningful representation of cell competition to predict the fate of cells. (a) A single cell in a mixed population of MDCK^WT^ (green) and scrib^kd^ (magenta) cells has two easily observable fates, mitosis or apoptosis. The central hypothesis is that the local tissue organization over time determines the fate. (b) Single-cell tracking is used to build a detailed training dataset of trajectories. The movies are truncated to remove images that encode the fate of the cell. The goal of the machine learning model is to learn a representation that can predict the fate of a cell (circled in white) given the local configuration of its neighborhood during interphase. Importantly, the model does not actually *observe* the fate, since recognizable morphological changes fall beyond the cutoff. Images are taken at 4 minute intervals, MDCK^WT^ cells appear in green and scrib^kd^ in magenta. (c) Overview of the *τ*-VAE model. The image data are encoded using a convolutional probabilistic encoder. The trajectory in feature space is the input of the temporal model. The output of the temporal model is a prediction of the future fate of the cell. During training, this can be compared to the known, ground truth, fate of the cell. (d) Training the probabilistic encoder using a *β*-VAE. The *β*-VAE is used to learn a low-dimensional representation of the image data. The cyan region represents the multivariate normal distribution of the image encoding. A sample is taken from this distribution to generate the synthesized image. (e) Using a large corpus of images (~1.2 million), we perform PCA on encodings in latent space to explain the variability of the dataset. The encoding can then be projected using these vectors to yield the features.

In this study, we sought to examine a new scientific paradigm – using Artificial Intelligence (AI) to uncover the determinants of cell fate directly from a large corpus of time-lapse microscopy data. Specifically, we sought to explore the possibility of learning an interpretable and predictive model of competition using an unbiased approach.

Recent studies have shown that machine learning (ML) is adept at uncovering complex patterns in microscopy data [10, 11]. In conventional feature engineering approaches, prior knowledge is incorporated into a model by choosing features that represent the system, for example, by measuring image properties or adding relevant fluorescent cell signalling reporters. This has recently been used, with ML-enabled dimensionality reduction, to study transitions in human pluripotent stem cell populations [12]. However, choosing appropriate measurements becomes increasingly difficult with more complex features such as describing the local organization of tissues comprising multiple cell types and varying degrees of epithelialization. One promising method is the use of unsupervised deep learning methods, such as variational autoencoders (VAE, [13]). A VAE learns a probabilistic approximation of the underlying distribution of data, meaning that the latent representation can be used as descriptive features of the system. Several recent studies have utilised autoencoders to encode complex cell shapes and other visual features in an interpretable manner [14, 15, 16, 17]. However, these studies have typically been performed on sparse, isolated cells and usually as single observations in time. Other models have attempted to explicitly incorporate time. For example, a recurrent neural network was used to predict lineage choices in hematopoietic stem cells [18]. However, this architecture does not lend itself to introspection and therefore does not directly provide any interpretable insight into the biology.

Here, we sought to learn a model of cell behaviour directly from time-lapse image data. We expand upon the use VAEs to encode cell shape and incorporate local tissue organization as well temporal features, to learn an explainable model of a complex, physiologically important biological phenomenon, cell competition. Finally, we introduce a novel discriminator network, that uses this learned model to identify drugs that affect the underlying mechanism of cell competition.

## Results and Discussion

### Data acquisition and training data

We used a well-described model of cell competition consisting of co-cultures of mammalian MDCK wild-type (MDCK^WT^) and a variant expressing an shRNA targeting the polarity protein *scribble* that can be induced to become mechanical loser cells by addition of tetracycline (scrib^kd^) [19]. To differentiate scrib^kd^ from their MDCK^WT^ counterparts, we expressed nuclear markers fused to different fluorescent proteins (e.g. H2B-GFP for MDCK^WT^, and H2B-RFP for scrib^kd^). Cells were seeded in different ratios, and then, using automated time-lapse microscopy we followed the evolution of the competition over periods of 80 h, taking images at 4 min intervals. We collected 111 independent movies, totaling 7,768 hours of competition experiments (**Fig 1a**).

From this dataset, we extracted single-cell trajectories, making sure that we could observe the entire lifespan of each cell including its fate (either mitosis or apoptosis, **Fig 1b**). To do this, we segmented the time-lapse image data using a fully convolutional residual U-Net [20], then used a dedicated convolutional neural network (CNN) to classify each nucleus into one of five states (interphase, prophase, metaphase, anaphase or apoptotic) based on image features [8]. Then, we tracked all cells over time [21]. Next, we classified the fate of each track as either mitotic, apoptotic or unknown using a dedicated cell fate classification network (**Extended Fig 1**, **Supplementary Information**). We discarded trajectories with an unknown fate. We manually verified all apoptotic trajectories and a subset of mitotic trajectories to confirm that the fate labels could be used as ground truth for the purposes of training a model. Together with the fact that two distinct cell types interact, this CNN encodes the entirety of the *a priori* scientific knowledge that we use to pose our problem. In total we acquired 36,062 mitotic and 2,225 apoptotic trajectories, distributed across the two cell types (**Supplementary Information**). Since we do not know *a priori* what information is required to predict cell fate, we extracted glimpses [22] which capture three different spatial scales of the neighbourhood surrounding the cell of interest (Small, Mid and Large, corresponding to 21×21 μm, 42×42 μm and 84×84 μm FOV respectively) but contain the same number of pixels (64 × 64 px). Further, for the mid-scale view, we also applied masking (using segmentation masks) to artificially remove either the central cell or the neighbors from the image data (**Extended Fig 2**) to test which representation was most salient. These five datasets could then be used to determine the best performing models and determine where the information necessary for fate prediction is contained.

### An explainable model for cell fate prediction from image sequences

Having acquired a large training set, we next designed a machine learning framework to learn the features of a single cell and its neighbourhood over time, which act as strong predictors of cell fate (**Fig 2a**). One of the design goals of this system was to have minimal human prior insight integrated into the model. First, we sought to learn an interpretable representation of the image data in an unsupervised manner. We trained a variational autoencoder (*β*-VAE, **Methods**, [13, 23]) to learn a compact latent representation of cell image data using 1.2 million different images of individual cells and their first neighbours (**Fig 1e**). The *β*-VAE learns a low dimensional representation of the image data using a probabilistic encoder. This low dimensional representation (bottleneck) should encode the image in a minimal number of parameters, in the same way that a human operator might describe the image in terms of the cell type, orientation of cells, local neighborhood and so forth. Next, a decoder attempts to reconstruct the real image using this low dimensional representation. The objective function guides the *β*-VAE to learn a representation of the image data which is expressive enough to reconstruct the original features, but where each of the latent dimensions are independent and interpretable. We modified the training objective to linearly increase the *capacity* of the bottleneck of the *β*-VAE during training. At low capacities the network is forced to prioritise matching the approximate posterior to the exact posterior distribution. At higher capacities, the network prioritises the quality of the reconstruction. During training the *β*-VAE first learns to represent gross level information such as cell type, before features such as cell shape and local organization (**Supplementary Information**).

**Figure 2:**
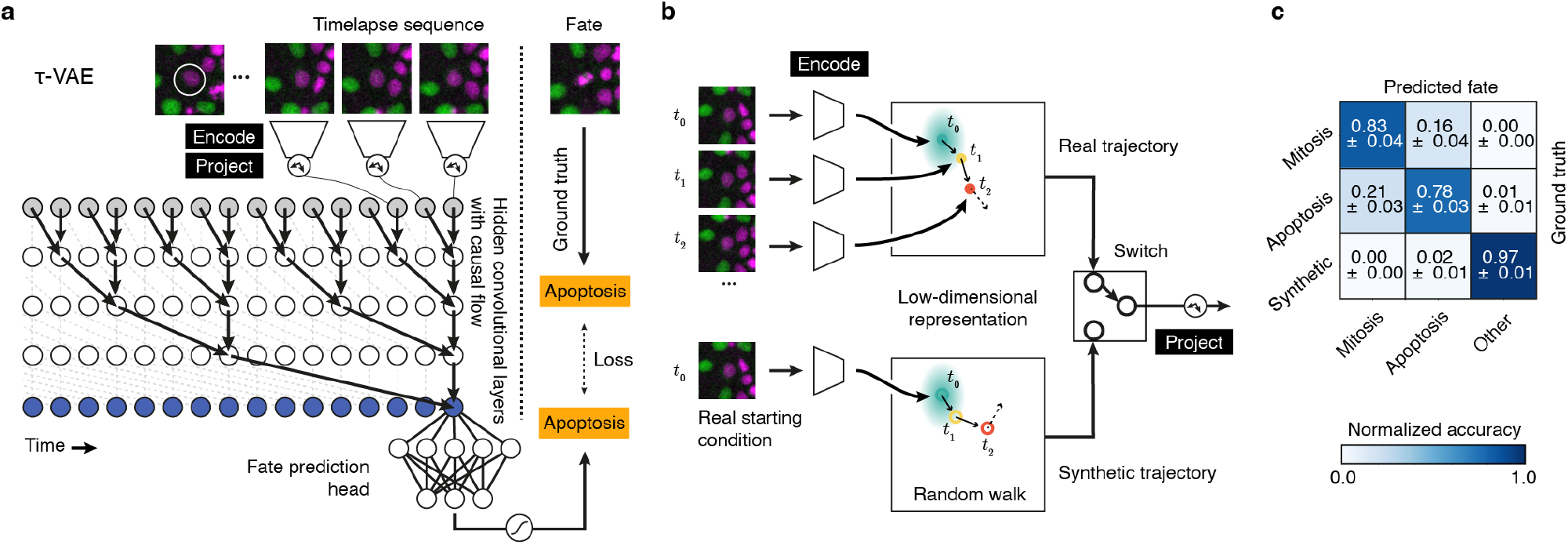
Architecture, training and inference using the *τ*-VAE. (a) Schematic of the *τ*-VAE architecture. A TCN network uses the probabilistic encoding, followed by projection of the trajectories to predict the fate. The TCN network has a receptive field of 128 time steps (~8.5 h prior to the event) and seven hidden layers consisting of 64 convolutional kernels. Only five layers are shown for simplicity. The fate prediction head has three output layers corresponding to “apoptosis”, “mitosis” and “other”. (b) During training of the model, mini-batches of data are created using a mixture of real and synthetic trajectories. Synthetic trajectories are generated *on-the-fly* during training, by encoding a single, randomly sampled image from a real trajectory, and using this encoding as a starting point for a random walk in latent space. A “switch” randomly chooses samples from either the real data or synthetic data to form the mini-batches during training. (c) Confusion matrix for best performing model on scrib^kd^ trajectories using randomly selected trajectories (*n* = 300) for testing across a 10-fold cross validation. Errors represent the SEM.

By encoding a large set of images we were able to perform Principal Component Analysis (PCA) in the latent space, which yields linear features that explain the variance in the entire dataset (**Fig 1e**). We use these principal components (PCs) as the inputs to the temporal model for making predictions (**Fig 2a**). Whilst this projection step is not required for correct prediction of cell fate, it enhances the interpretability of the learnt features (see discussion).

Next, we built a prediction network that utilises the temporal sequence of images of each single-cell trajectory to output cell fate. Based on current biological knowledge, we reasoned that the biochemical commitment to apoptosis or division occurs hours before we observe the fate, so we trimmed each trajectory to remove observations that show recognizable morphological features of mitosis (such as DNA condensation in prometaphase and alignment of chromosomes during metaphase) or apoptosis (such as DNA fragmentation) (**Supplementary Information**). As such, the prediction network is forced to use only features from the time evolution of the interphase cell and its neighbours to make a prediction about the fate.

Our prediction network (τ-VAE, **Fig 2a**) couples a convolutional probabilistic encoder and projector to a temporal convolution network (TCN, [24, 25]) to enable cell fate prediction within an explainable framework. The TCN uses *causal convolutions* to ensure the prediction at time *t* is only dependent on the observations [*x_t–r_* ··· *x_t_*], where *r* is the receptive field (equivalent to a window of time before an event, ~8.5 h in this case) of the TCN (**Supplementary Information**). The output of the TCN is connected to a dedicated prediction head, a densely connected network with three outputs corresponding to “apoptosis”, “mitosis” and “other”. A final softmax activation yields the prediction of the full network. Overall, The TCN takes a sequence of observations of a single-cell in interphase and returns a prediction for the cell fate, without ever observing the fate (**Fig 2a**). In training the *τ*-VAE, we supplemented the real data corresponding to mitotic and apoptotic classes, with a dynamically generated “synthetic” class to simulate trajectories which were neither apoptotic nor mitotic, corresponding to the “other” output class (**Fig 2b, Extended Fig 3, Methods, Supplementary Information**). This is important as we do not want the model to learn only the features of mitosis, and predict apoptosis by exclusion, or *vice versa*. Further, these synthetic trajectories, while exploring a similar region of feature space, have no causal relationship with cell fate. In adding these synthetic trajectories, we ensure that the model learns the features predictive of both apoptotic and mitotic events.

We tested the five different representations of the image data (**Extended Fig 1**) as input to the prediction network. In all representations, the image data consisted of the same number of pixels, but comprised either different spatial scales or lacked information about either the central cell or the neighbors. We trained the *τ*-VAE network and measured the accuracy by comparing the predicted fate with the ground truth fate on a set of unseen test data consisting of 300 trajectories. For each dataset, we split the data by the cell type (MDCK^WT^ or scrib^kd^) and calculated separate confusion matrices to ensure that there was no systematic bias in the predictions. To account for any potential bias in the testing set, we performed *k*-fold cross-validation (*k* = 10) on each model (**Methods**), randomly choosing a subset of training and testing data in each validation. Overall, the best performing networks used only the central cell region ((“Small View, All Cells” model) to predict the fate, with an average fate prediction *F*_1_-score of 0.87 ± 0.02, across both cell types as shown in the confusion plots (**Fig 2c**, **Supplementary Information**). For reference, a TCN trained using a set of human chosen image features (those shown in **Fig 3a**) achieves a lower *F*_1_-score of 0.71, and is particularly poor at identifying apoptosis, yielding an accuracy of 0.49. This demonstrates that the *β*-VAE is able to capture more salient image features, enabling a more accurate fate prediction, by learning them directly from the distribution of the data.

**Figure 3:**
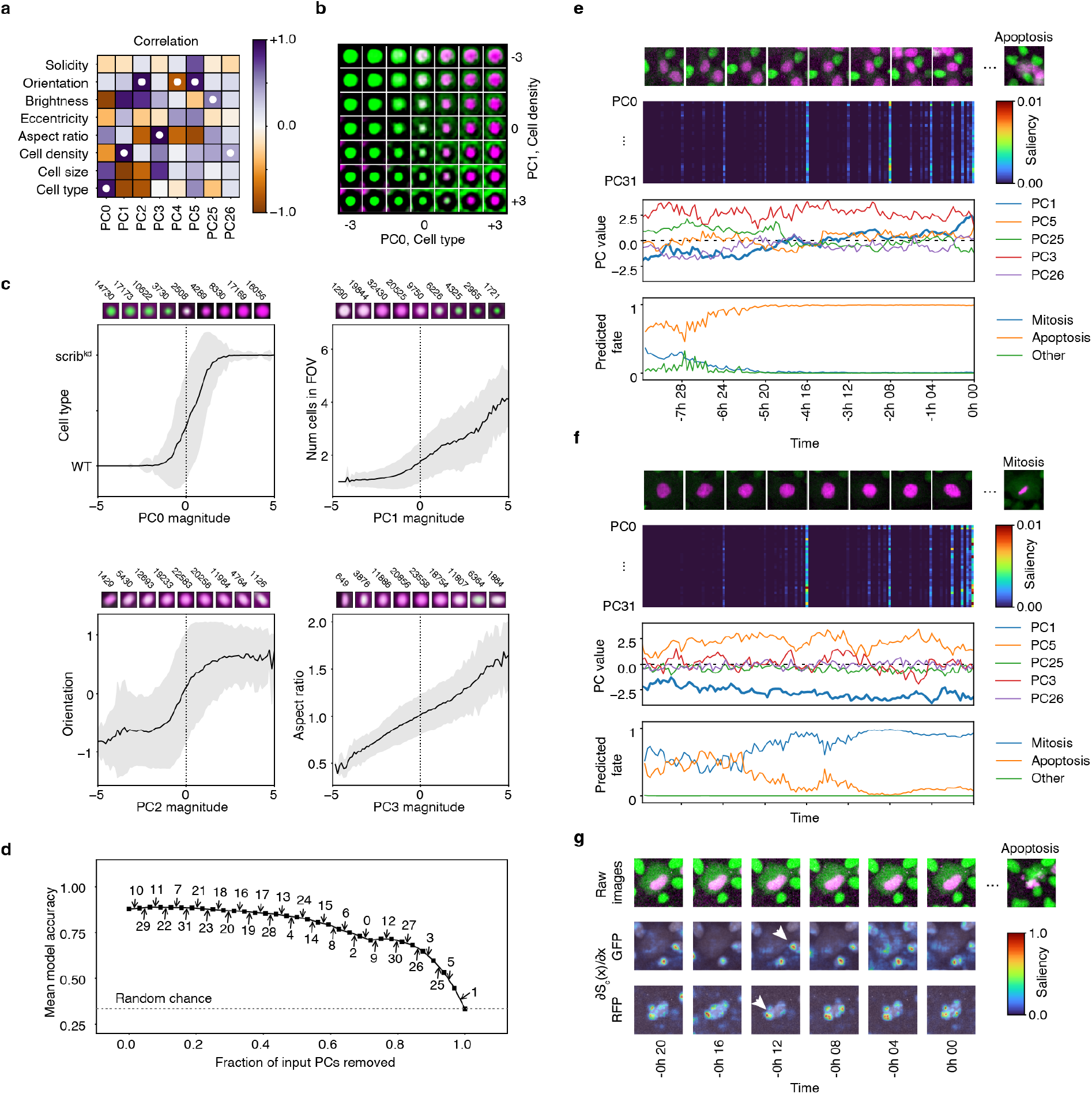
An explainable internal representation of cell competition. (a) Pearson correlation coefficient between principal components and measurable physical parameters. White dots denote the parameter with the highest correlation to each PC. (b) Linear combinations of PC0 (cell type) and PC1 (cell density) over the range [-3, 3]. (c) Interpretability of the projected *β*-VAE latent space, showing correlation between PC0-3 and known physical parameters. Continuously varying the PC changes the state of the cell in an explainable manner. Above each plot are average images with binned encodings of the respective PC value. Numbers represent the distribution of these states in the dataset, calculated from a sample of images (*n* = 100, 000). Errors represent SD. (d) Feature ablation demonstrates the role of each PC in the final prediction of the model. Each arrow indicates the cumulative replacement of a given PC with Gaussian noise. Error bars represent the standard deviation of model accuracy over all ten TCN models. (e) Example apoptotic trajectory with network predictions, and internal representation. Top row shows a sampled sequence of images from the trajectory. Second row shows the PC feature saliency over time calculated by the TCN. Third row is the values of the top-5 principal components over time. Final row is the prediction of cell fate over time. (f) Example mitotic trajectory with network predictions, and measured parameters, as in panel e. (g) Feature saliency *w.r.t*. the input pixel data obtained by backpropagating error through the *τ*-VAE. The middle and bottom rows show the normalized pixel saliency (*∂S_c_*(*x*)/*∂x*) in the GFP and RFP channels of the input (top row), respectively. White arrows indicate examples of regions of high saliency corresponding to nearby dividing MDCK^WT^ cells or changes in the nuclear morphology of the target cell.

The TCN was preferred over other time-series models, such as a Recurrent Neural Network (RNN), primarily because it demonstrates a longer memory than recurrent architectures with the same capacity and have been shown to outperform RNNs [24, 25]. Further, the TCN was deemed to possess a higher degree of interpretability. The TCN works similarly to a convolutional neural network; it performs feature extraction on different regions of the input, then pools lower-level features into higher-level features progressively until a classification is reached (**Methods**). The fact that a TCN processes all time-steps of the input simultaneously enables the determination of which regions of the input (either, which time-steps or which features) were most responsible for a particular classification. Inspection of this kind is much more difficult when using RNNs, where each time-step of the input is processed sequentially and separately. Moreover, when we trained and tested a Long-Short Term Memory (LSTM [26]) RNN on the same scrib^kd^and MDCK^WT^data – using the same procedures – we found that it predicted cell fate with lower accuracy (*F*_1_-score of 0.82 ± 0.04, **Supplementary Information**), compared to a TCN model that possessed a similar number of trainable parameters (*F*_1_-score of 0.87 ± 0.02, **Supplementary Information**). The TCN contained 123,302 trainable parameters, whereas the LSTM contained 136,195 trainable parameters. Notably, the LSTM network performed worse on the prediction of apoptosis (recall of 0.71 ± 0.03) than the equivalent TCN backbone (recall of 0.78 ± 0.03).

We used the best performing *τ*-VAE network (“Small View, All Cells” model, *i.e*. cropped to the central cell at the highest magnification) for all further analyses. We concluded that the *τ*-VAE network is able to accurately predict the cell fate based on the interphase local tissue organization alone, having learnt features directly from the image data to enable this task. Next, we sought to introspect the model and to assign meaningful semantic labels to the learnt features.

### Interpreting the model

The goal of the *τ*-VAE network is to learn an end-to-end model that requires minimal input from experts to predict the fate of cells in competition. The implicit hypothesis is that there is sufficient information in the observations of local tissue organization to enable this prediction. Having determined that our approach is able to accurately predict the fate of cells in a competitive system, we sought to interrogate the learnt features of the *τ*-VAE. In contrast to other approaches, we do not perform feature engineering to select parameters that define the problem (such as cell density, number of neighbours, etc), but rather, we extract these automatically and directly from the data, based on the latent representation of the *τ*-VAE. We used several different approaches to interpret the model.

#### Assigning physical parameters to the latent features

Since the training objective of the *β*-VAE encourages a continuous, but disentangled internal representation of the image data, we sought to assign meaningful semantic labels to those latent variables. Analysis of the latent features revealed that some parameters show covariance, an insight that can be confirmed by direct examination of the image data – for example, in the dataset, there is a correlation between cell type and nuclear area, since scrib^kd^cells tend to have larger nuclei than MDCK^WT^cells (**Supplementary Information**). To remove covariance, we performed PCA on the latent space (**z** ∈ ℝ^32^) yielding 32 principal components ordered by the magnitude of the variance explained by the component. We analysed the correlation of the components with parameters that could be measured from the images (**Fig 3a**). Inspecting these components shows that the first two (PC0 and PC1) account for 26.9% of the variability of the data, and seem to represent cell type and cell density (number of cells visible in the glimpse) respectively (**Fig 3a-c**, **Extended Fig 3**). Component PC2 encodes an orientation parameter of the central nucleus, while component PC3 encodes nuclear aspect ratio (**Fig 3c**). Higher principal components, like PC25 encode parameters such as fluorescence intensity of H2B-GFP/RFP but the correlation coefficient is weaker. We confirm these assignments by sampling images from the dataset with various values for these components (**Supplementary Information**). Later components broadly enable the network to encode the arrangement and identity of cells in the local neighbourhood (**Supplementary Information**). Strikingly, projecting the *β*-VAE latent space enabled us to learn an explainable model of the local tissue organization of cells in a completely unsupervised manner. Next, we sought to investigate the role of these principal components over time in the prediction of cell fate.

#### Feature ablation studies to determine the minimal information required for prediction

To determine the minimal information required for cell fate prediction, we removed individual principal components in a systematic manner (replacing them with Gaussian noise at all time steps) and calculated the performance of the network after each component removal. Ablated networks were ranked according to their effect on the prediction accuracy. Through multiple iterations, we found that a single component (PC1 - nuclear area/cell density) could be used to predict cell fate with 43 ± 2% accuracy – significantly higher than random chance assuming an equal probability of choosing any fate (33.33%, **Fig 3d**). In the ablation approach, PC1 was the last component to be removed, suggesting the single highest contribution to the prediction accuracy. The ablation study reveals that the top five components (PC1, PC5, PC26, PC3, PC26) account for 64% of the prediction accuracy, with the remaining 27 components contributing a further 36%. Importantly, when all inputs are replaced by noise, all fates are predicted with equal probability, suggesting no inherent bias toward any fate in the network. Remarkably, this suggests that, in line with our current understanding of mechanical cell competition stemming from nearly a decade of experimental studies [19, 7, 27, 8, 9], our model has autonomously learnt that cell density is a strong predictor of cell fate, directly from the data with no expert input.

#### Timescales of predictions and feature saliency to visualize network attention

Since the TCN is able to output a prediction after every additional time step, we can use the prediction head (**Fig 2a**) to visualize the time evolution of the prediction (**Fig 3e-f**). Across all of the data, we find that for the scrib^kd^cells, the *τ*-VAE network predicts apoptosis early (up to 8h before an event) and mitosis late (~2h before an event, **Supplementary Information**). These long timescales suggest that the predictive power of our model is not due to previously unrecognised changes in nuclear morphology that would be indicative of apoptosis or division. Further, we can inspect the magnitude and contribution of each principal component to the prediction. In general, we found that PC1 (related to cell density) was often much larger in those trajectories undergoing apoptosis (**Fig 3e-f**, **Supplementary Information**). To assess the model’s use of components to make predictions, we utilise the error gradients during backpropagation in the network to calculate the saliency (**Methods**, [28]). Feature saliency is a method to determine which features of the input contribute most significantly to the classification accuracy of the *τ*-VAE. We computed this in two ways (i) PC feature saliency, *i.e*. the PC input to the TCN network (**Fig 3e-f**) and (ii) raw pixel saliency *i.e*. the raw pixel input to the encoder network (**Fig 3g**, **Supplementary Information**). The latter approach identifies the raw image pixels that contribute most to the eventual prediction. The PC feature saliency reveals the timescales of activations within the network that are used to make predictions. The high gradients reveal that features throughout the timeseries are used in making the predictions for apoptosis and mitosis (**Fig 3e-f**). Empirically, the pixel saliency reveals that nearby cells and the geometry (aspect ratio, convexity) of the nucleus (**Fig 3g**, **Supplementary Information**) have significant contributions to the prediction - however, it is difficult to assign quantitative meaning to these observations, owing to the fact that they are in image space rather than feature space.

### Challenging the learned representation with biochemical perturbations

Finally, to confirm that the network has learnt a model of mechanical cell competition, we sought to challenge the model with cells treated with different biochemical perturbations. For example, we performed experiments using cells that were uninduced (MDCK^WT^:scrib^kd,tet-^) such that there was no competition. In this case, the knockdown of the polarity protein Scribble is not induced with tetracycline (scrib^kd,tet-^), so the cells do not engage in mechanical competition with MDCK^WT^ cells, but can still be distinguished by their H2B-RFP marker. We acquired timelapse data of the cells and confirmed that the scrib^kd,tet-^ cell count was consistent with a non-competitive scenario (**Fig 4a**, **Supplementary Information**). From this dataset, we randomly selected a set of full length trajectories with a known fate (either apoptosis or mitosis), and, after removing the final images and discarding the trajectories where no event occurs, we passed them to the *τ*-VAE network. We found that the fate prediction accuracy of the network dropped significantly for the scrib^kd, tet-^ cells. Strikingly, many scrib^kd, tet-^ mitotic trajectories were predicted to be apoptotic as can been seen from the confusion matrix (**Fig 4b**). This is an important result – the τ-VAE, using the available information predicts, correctly, that the scrib^kd, tet-^ cells should die under these conditions if knock-down of Scribble had been induced to start competition. The *τ*-VAE predicts an outcome that is consistent with its model of the phenomenon, based on the data available. A human would arrive at the same conclusion, given the same information. The fact that, under an unseen biochemical perturbation that disturbs the competition, we subsequently observe that H2B-RFP marked cells do not die, lays the foundation for a method to identify systematic deviations from the normal behavior that the model was trained to predict. Furthermore, this discrepancy indicates that the model is predictive, and has learned the determinants of cell fate *specific to* cell competition, rather than relying on previously unrecognised changes in central cell nuclear morphology. Indeed, if an equal performance on the scrib^kd, tet-^ datasets had been observed, this would have implied that the model was simply classifying image features indicative of mitosis or apoptosis in the observed cell.

**Figure 4:**
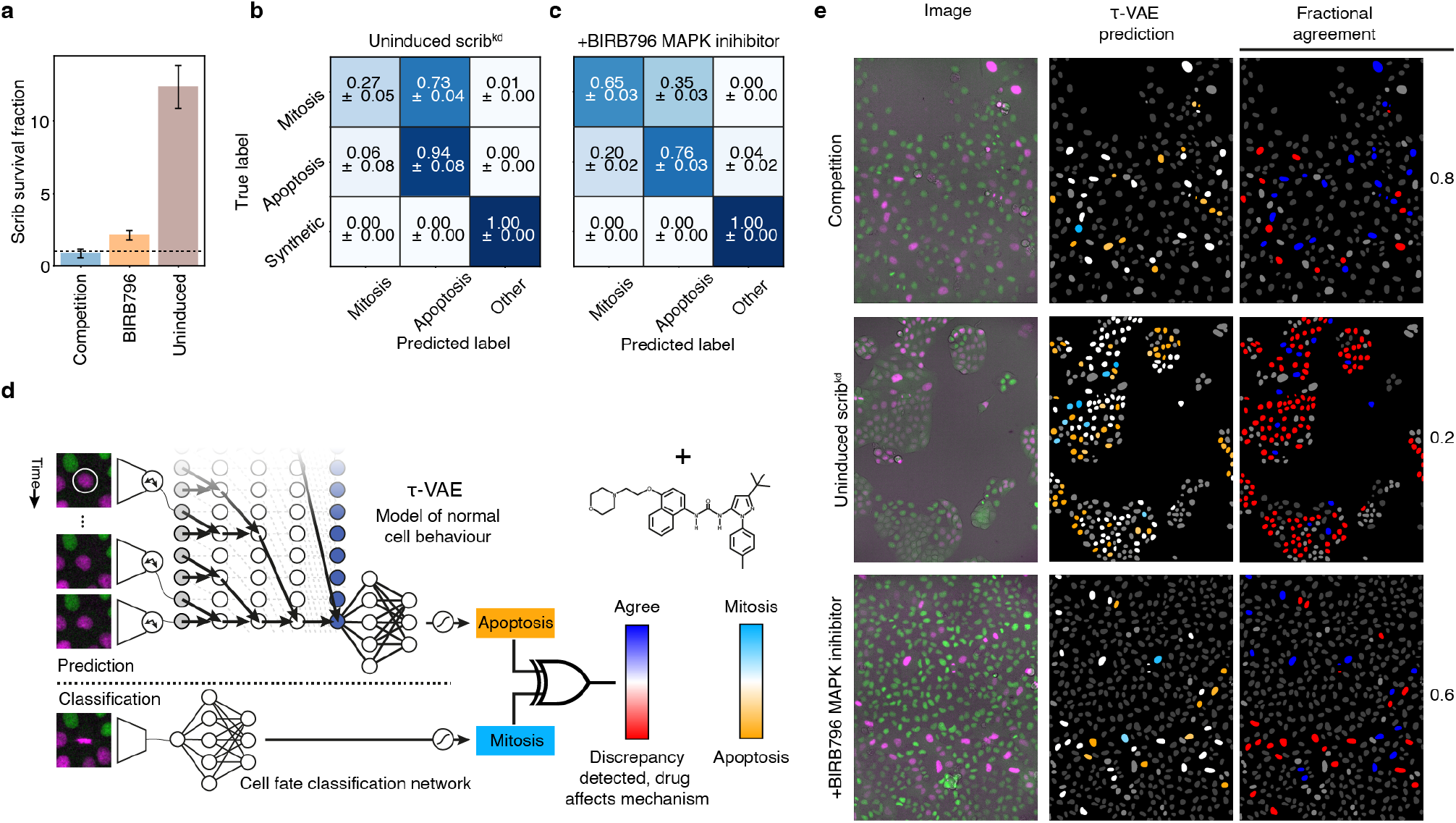
The predictive model of cell behavior enables drug evaluation. (a) Survival fraction for scrib^kd^ cells in competition (MDCK^WT^:scrib^kd^), BIRB796 treated (MDCK^WT^:scrib^kd^ + 2 μM BIRB796) and uninduced (MDCK^WT^:scrib^kd,tet-^). Values below 1 (dotted line) indicate a cell population decrease, values greater than 1 indicate a population increase over the course of the experiment. (b) Confusion matrix of prediction accuracy for scrib^kd,tet-^ cells (*n* = 161 real trajectories) showing many mitoses are incorrectly predicted as apoptoses. (c) Confusion matrix for BIRB796 treated scrib^kd^ cells (*n* = 198 real trajectories) showing a similar pattern to the scrib^kd, tet-^ condition. (d) A discriminator network that uses two models to detect changes in cell behavior. Network A is the *τ*-VAE model of learned cell behaviour and Network B is the cell fate classification network. A discriminator (shown as an XOR gate) determines discrepancies between the two outputs. (e) Example outputs of per-cell predictions and discriminator output for individual timepoints of competition, uninduced and BIRB796 treated timelapse movies. The left column shows the raw input image data. The middle column shows the current prediction (in this frame of the movie) of the *τ*-VAE network for scrib^kd^ cells in the FOV. Blue represents mitosis, orange represents apoptosis and white represents unknown fate or insufficient data to make a prediction at this time-point. The right column shows the discriminator output for each cell in the FOV. Red indicates that the *τ*-VAE network did not agree with the fate classification, and blue indicates agreement. Grey cells have no predictions associated with them as they are MDCK^WT^ cells.

This variation in the performance of the *τ*-VAE when applied to cells under different biochemical perturbations suggested that our model is sensitive to changes in gene expression and the biochemical mechanisms of competition. Therefore, we sought to determine whether the methodology could be used for identifying drugs that perturb competition without further modification to the prediction network. Recent studies have suggested that p38 kinase inhibitors may interfere with mechanical cell competition by inhibiting the stress response pathways that lead to apoptosis [7]. To test this hypothesis, and to determine whether the network was able to detect treatments that perturbed competition, we acquired timelapse data of the cell competition (MDCK^WT^:scrib^kd^) in the presence of 2 μM BIRB796, a p38 MAPK inhibitor [29]. We measured the loser cell count over the course of the experiments and noted that there was a higher survival fraction of the scrib^kd^ cells, although they still grew significantly slower than MDCK^WT^ (**Fig 4a**, **Supplementary Information**). From these data, we extracted single cell trajectories and used the *τ*-VAE network to predict the fate of the cells as before. As with the uninduced dataset, the network predicted a significantly higher number of apoptoses where the true label was mitosis (**Fig 4c**). Indeed, the τ-VAE network predicts which cells should have died if stress response pathways had not been inhibited.

Given that both the p38 MAPK inhibitor and the scrib^kd,tet-^ condition interfere with the competition by limiting apoptosis, we would expect the scrib^kd^ cells to reach higher densities in these conditions. Indeed, when analysing the network’s representation, we noticed that the signatures of incorrectly predicted trajectories are more similar to the trajectories categorised as apoptotic under control conditions, especially with respect to the increased magnitude of PC1, that represents local cell density (**Supplementary Information**). This is consistent with the observations that the scrib^kd, tet-^and BIRB796 treated scrib^kd^cells reach higher densities, with significantly lower apoptotic rates. Overall, our results suggest that the *τ*-VAE network, trained on the MDCK^WT^:scrib^kd^ data, has learnt a complex and predictive model of cell competition, that is sensitive to local changes in the tissue organization and the signalling pathways participating in competition.

#### A method for automated drug screening

Having established that the *τ*-VAE network is able to represent a complex model of cell behavior in an explainable manner, we sought to define a general approach to utilise such a predictive model for image-based drug screening [30]. To do so, we introduce a novel discriminator network (**Fig 4d**) that compares the outputs of two models (Networks A & B) to determine discrepancies indicative of drug activity or other perturbations of the molecular mechanisms. The two networks utilise different amounts of information from each single-cell trajectory. Network A (the *τ*-VAE), a model of “normal” cellular behavior during cell competition, uses only the early part of the trajectory to predict the fate of the cell. In contrast, Network B (the cell fate classification network, **Supplementary Information**) uses the entire trajectory to classify the actual fate of the cell. When these two models agree for a particular cell, it suggests that the fate is predictable and thus competition conforms to learned behavior. However, when the two models disagree, this suggests that the cell is deviating from normal learned behavior, presumably due to the influence of an added drug or other perturbation. We demonstrate the utility of the discriminator to evaluate individual scrib^kd^ cells in movies in three different conditions, MDCK^WT^:scrib^kd^, MDCK^WT^:scrib^kd, tet-^ and MDCK^WT^:scrib^kd^ + 2 μM BIRB796. First, we determined the performance of the cell fate classification network, finding it to achieve an accuracy of 0.96 in determining cell fate (*n* = 392) across all of the data. Then we used the *τ*-VAE to make predictions for these cells, and the discriminator to determine the agreement with the cell fate classification network. In control conditions, the two networks show a high level of agreement with a fraction of agreement *a* of 0.8. In contrast, in the presence of a drug or other perturbation, the two networks disagree. The uninduced Scribble (*a*=0.22) and and BIRB796 (*a*=0.62) treated cells show a lower fraction of agreement between the *τ*-VAE and the cell fate classification, indicating deviation from the normal model of behavior (**Fig 4e**). Therefore, the discriminator network automatically identifies conditions that deviate from learned behavior. As an example, our approach could be used to identify signalling pathways acting during competition by screening libraries of siRNA or chemical compounds whose targets are already known. In this case, we would expect that our discriminator should show high perturbation when we target the proteins forming part of signalling cascades involving in mediating the detection and outcome of competition.

## Conclusions

Deep learning is now a powerful tool in microscopy image analysis [10, 11]. However, the complex internal representations of many deep learning models, and the difficulty of analysing time dependent features, means that they have rarely been used to gain mechanistic insight into biological phenomena. Here, we developed an end-to-end machine learning model capable of discovering the physical parameters and rules of a complex biological phenomenon, cell competition, directly from image data. Starting *tabula rasa*, we demonstrate that our approach is able to learn a meaningful representation of cell behaviour in an automated and unbiased manner. Strikingly, the model learns that, local cell density is the single most important determinant of cell fate in mechanical competition, an observation that has taken scientists considerable experimental research and data analysis effort to determine. Most exciting is that we are able to introspect the model to identify the physical features enabling prediction as well as the time-scale over which correct predictions are made.

Many design decisions involved in building such a model require input from human experts. These include how the scientific question is posed as a machine learning problem, the choice of input data, the model architecture and the choice of hyperparameters used to train the model. We discuss some of these below.

First, we pose the hypothesis that cell fate in competition is determined by the local tissue organization, as a machine learning problem. If a causal link between the local tissue organization over time and the fate of a cell exists, then a ML model should be able to predict the fate of the cell given a sequence of images. Therefore, an implicit assumption is that the image data encode all of the information required to make a prediction about the future state of a cell. Remarkably, our *τ*-VAE model, using the data provided, successfully identifies local cell density as a strong predictor of single-cell fate in mechanical competition. We expect that the interpretability of principal components will prove invaluable in formulating hypotheses about the nature of the mechanical changes governing competition and the signalling pathways leading to loser cell death.

Furthermore, the ability of the model to *predict* cell fate ahead of time, rather than merely *observe* early indicators such as morphological change, is evidenced by the fact that the model showed discrepancies between the predicted and observed fates when applied to biochemically perturbed datasets (**Fig 2e-f**). If the model was simply observing previously unrecognised early indicators of cell fate, it should have performed as well on the perturbed datasets as on the control datasets.

An extension of this work would be to use the same approach to identify how density regulates fate during competition by providing the *τ*-VAE with additional information. For example, the addition of the temporal evolution of junctional tensions determined through Bayesian inference or timelapse data of live fluorescent reporters for known biochemical pathways could be added as an additional data sources. Indeed, previous work has shown that an early event leading to loser cell death is the activation of stress pathways such as p53, p38, and JNK upstream of caspase-3 and apoptosis [7, 31]. By introducing reporters for these pathways, it may be possible to learn patterns of biochemical activity that are predictive of cell fate, yielding insight into how fate is regulated and how these stress pathways combine in time in the decision to apoptose.

Another critical decision is the choice of model hyper-parameters, such as spatial scale, which can emphasize cell-autonomous or cell-cell interaction features. This is difficult to achieve using feature engineering since defining simple metrics that capture local tissue organization is extremely challenging. In our approach, the spatial scale of glimpses emphasizes different features of cell or tissue organization and allows the encoder to determine interpretable features that best represent the system. In an unbiased assessment of information arising from different scales, we found that much of the salient information regarding local tissue organization could be encoded in the smallest scale. For example, changes in local cell density were reflected in a compaction of the central cell nucleus and encroachment of neighboring cells into the glimpse, and encoded in a single component of the learned representation (as can be seen in **Fig 3b**).

In building machine learning models to augment scientific research, we are confronted by a trade-off between predictive power and explainability in the design. Forcing a model to be explainable may actually reduce its predictive power, since the representation may not encode the data as efficiently. Recent models such as vision transformers [32] may prove interesting architectures for future work. However, beyond simple visualization of network attention, the interpretability and quantification of image data using such models currently seems challenging [33]. Another interesting avenue may be to utilize geometric learning [34] to represent each cell in a tissue as a node in an evolving graph. In our *τ*-VAE approach we have sought to balance explainability with prediction accuracy for cell fate.

Overall, our *τ*-VAE approach requires minimal human input to train, is able to correctly predict the fate of cells in mechanical cell competition, and learns an interpretable representation of the behavior. None of the previously-reported results regarding the determinants of cell fate in mechanical cell competition were used to assist the model in learning the behavior of the cells that it observed; the *τ*-VAE learnt these rules autonomously – *i.e*. without specialist knowledge of cell competition. Although we have demonstrated the utility of this approach using a model phenomenon and cell type, the approach is generalizable to many other systems, for example the study of the micro-environmental factors leading to differentiation of stem cells or embryonic development.

Finally, our *τ*-VAE model can be used directly in combination with high-throughput screening approaches, and with no further modification, to investigate which signaling pathways participate in competition by screening libraries of drugs with known targets or siRNA. We have shown that a novel discriminator network, based on the learned model, can detect conditions which deviate from the normal cellular behaviour due to biochemical inhibition or silencing, signifying that it can be used as a screening tool. Once trained, this fully automated system is able to discriminate between normal cell behavior and perturbations without any further human intervention, paving the way for mechanism-aware AI-based drug discovery.

## Methods

### Cell culture, imaging assays, single-cell tracking and cell fate classification network

Detailed methods can be found in the supplementary information.

### Drug treatments

BIRB796 (Tocris 5989, [29]) was dissolved in DMSO and added to the cell culture 5 h before imaging at a final concentration of 2 μM. As a negative control, a similar volume of DMSO was added to some wells.

### Variational Autoencoder (*β*-VAE)

We built a convolutional variational autoencoder to learn a low-dimensional representation of the cell image data that could be used by a TCN for fate prediction. The encoder network consists of four convolutional layers with 3 × 3 kernels, Swish activations, with 32, 64, 128 and 256 kernels respectively. Each layer was pooled by a 2 × 2 max-pooling operation. The convolutional output was flattened and split into two arms with two fully connected layers for the *μ* and *σ*^2^ estimators. We found that adding additional fully connected layers of 128 units between the estimators and the flattened encoder output improved the model performance. We used a random normal sampler to generate samples from the distribution. The decoder is inverse of the encoder, using nearest neighbor upsampling between the convolutional layers.

We use the revised objective function [23, 35] to train the network:

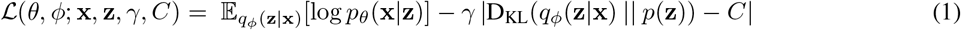

Our loss function is composed of two terms. The first term is the reconstruction loss term which penalises differences between the reconstruction (the decoded, encoded input) and the real input. In practice, we use the mean squared error between the input image (x) and the output (x′) of the *β*-VAE. The second term is the Kullback-Leibler divergence (D_KL_(· ║ ·)), which penalizes the latent space model variance from dropping to zero, by forcing the encoding to match the Gaussian prior with a diagonal covariance matrix, *N*(0, *I*). This has the effect of regularizing the latent space to promote a continuous representation of the underlying image data.

In our implementation, we dynamically adjust the bottleneck capacity (*C*) of the network during training. The value of *C* is scaled linearly as a function of training iteration, reaching a maximum value *C*_max_. This ensures that at early training iterations the network prioritizes the encoding, while at later iterations this is refined to optimize the decoding. The scaling constant *γ* balances the two terms of the loss function.

We prepared a training set of 1.2 million images of cells by sampling individual cells from the time-lapse movies. A fraction of cells (random 10% of cells in frame) was selected from a random sample of frames (10% of frames in movie) and an ROI of varying size around each cell was extracted. These were then downsampled using nearest neighbor sampling, to the network input size of (64 × 64 × 2). We then trained the *β*-VAE network using *γ* = 1000, *C*_max_ = 50, latent vector **z** ∈ ℝ^*n*^. We found *n* = 32 to be the optimal value of the latent space dimensions based on the reconstruction error and the explained variance in the dataset (**Supplementary Information**). Optimization was performed using an Adam optimizer for 100 epochs and a minibatch size of 256. Training images were augmented *on-the-fly* by random flipping, rotations and simulated edge cropping.

### Trajectory Synthesis

In order to simulate a third class of trajectory, referred to as “synthetic”, we utilised the generative property of the decoder network. The first frame of each synthetic trajectory is a real image, i.e. it is decoded by the decoder network from the latent-space encoding of an image that actually exists. For the next frame, the encoding is adjusted by adding to each latent variable a scalar that is sampled from a Gaussian distribution (with *μ* = 0.0 and *σ* = 0.2). The image for this second frame is then the decoder output with this new encoding as input. This process is then repeated until all the frames of the synthetic movie are generated, with the trajectories taking a random walk in latent space.

### Principal Component Analysis (PCA)

We used PCA to analyse the learnt representation of the total dataset. The principal components of the latent features derived by the *β*-VAE have a higher degree of interpretability than the latent features themselves. PCA was applied to the latent features before analysis of the latent space was undertaken. We used the *PCA* function from *Scikit-learn* to perform the decomposition into 32 components, once for each of the *β*-VAE latent dimensions.

### Temporal Convolution Network (TCN)

The timelapse sequences encoded by the *β*-VAE become the input features (*n* × *t*) of a temporal convolution network (TCN) [24, 25]. The procedure for preparing a timelapse movie for input to the TCN is as follows. First, each frame in the timelapse movie is normalized such that the pixel values of that individual frame have zero mean and unit variance. This is performed on a per-channel basis, such that the RFP and GFP channels are normalized separately. Next, the normalized timelapse movie is fed through the encoder network of the trained *β*-VAE. The encoder yields three outputs for each latent variable - the mean, standard deviation, and Gaussian-sampled value. This is calculated for every timelapse movie in the dataset.

Once encoded, the timelapse sequences are transformed from latent space (**Z**) into principal component space (**T**) using the transform **T** = **ZW**, where **W** are the principal components. and then fed into the TCN for training. The TCN employs a 1-D fully-convolutional network architecture, where each hidden layer is of the same length as the input layer [36]. To establish the “causal” nature of the convolutions, the architecture is set up such that a convolution output at time *t* is convolved only with elements at time *t* and earlier in the sequence, determined by the dilation factor [25]. The TCN is formed of seven stacked convolutional layers with respective dilations of 1, 2, 4, 8, 16, 32 and 64. Each of these layers has 64 convolutional filters. The output from the convolutional layers is fed into a fate prediction head. This network projects the output into a classification, and is composed of a fully connected layer with 128 units, and a final fully connected layer with three units that represent the possible classifications of the sequence (“apoptosis”, “mitosis” or “synthetic”). We used a sparse categorical cross entropy loss function to train the network.

The TCN is trained with a batch size of 128 for 100 epochs. Optimization was performed using the RMSprop optimizer with a learning rate of 0.001. Training sequences were augmented *on-the-fly* by random frame removal, cropping and the addition of noise to the encoded sequences. Regularization was ensured by applying batch normalization and a dropout rate of 0.3 to the TCN layer of the network.

### Feature saliency

We can determine the salient features of the input images by calculating a saliency map (*M_c_*) for an input (*x*). This is achieved by calculating the gradient of the output function *S_c_* with respect to the input during backpropagation: *M_c_*(*x*) = *∂S_c_*(*x*)/*∂x*. In practice, we average *n* gradient calculations, each with a small added Gaussian noise (*μ*=0.0, *σ* = 0.1 × |(max(*S_c_*) – min(*S_c_*)|) component to the input.

## Software availability

A reference implementation of the *τ*-VAE is available at: https://github.com/lowe-lab-ucl/cellx-predict [37]. For other enquiries contact the corresponding author.

## Data availability

Data is available from the UCL data repository: https://dx.doi.org/10.5522/04/16578959 [38]. For other enquiries contact the corresponding author.

## Author contributions

ARL and GC conceived and designed the research. GV performed experiments. CJS developed and performed computational analysis. ARL wrote the image processing and cell tracking code. CJS, GV, GC and ARL evaluated the results and wrote the paper.

## Acknowledgments

This work was supported by a BBSRC LIDo AI PhD studentship to CJS. GV was supported by BBSRC grant BB/S009329/1. We thank Nathan Day, Jasmine Michalowska and Dan Smaje for help annotating data, and Manasi Kelkar for additional supporting data. We also thank members of the Lowe and Charras labs for discussions and technical support during the project. ARL wishes to acknowledge the Turing Fellowship from the Alan Turing Institute. ARL and GC wish to acknowledge the support of BBSRC grant BB/S009329/1.

