## Supplemental Data 9 for "Learning the Rules of Cell Competition without Prior Scientific Knowledge"

### EXTENDED FIGURES

**Alan R. Lowe**[1,2,4,5*]

[1]Institute for the Physics of Living Systems
[2]Department of Structural and Molecular Biology
[3]Department of Cell and Developmental Biology
[4]London Centre for Nanotechnology
University College London, Gower St, London, WC1E 6BT, UK

[5]The Alan Turing Institute
Euston Rd, London NW1 2DB, UK

February 23, 2022

#### List of Figures

E1 **Data flow in the model.** (a) Example time-lapse microscopy data showing a mixed population of MDCK$^{WT}$ (green) and scrib$^{kd}$ (magenta) cells. (b) Single-cell tracking is used to build a detailed training dataset of trajectories. The single-cell track is used to extract a glimpse of the cell over time, that becomes the input data for the machine learning models. (c) The data preparation and inference pipeline. A CNN/LSTM network classifies the fate of the cell and determines the cutoff point to truncate the track to remove images that encode the fate of the cell. The goal of the machine learning model is then to learn a representation that can predict the fate of a cell (circled in white) given the local configuration during interphase. Importantly, the model does not actually *observe* the fate, since these data fall beyond the cutoff. Images are taken at 4 minute intervals, MDCK$^{WT}$ cells appear in green and scrib$^{kd}$ in magenta . . . . . . . . . . 2

E2 **Glimpse extraction and cell masking to determine the best image input scale for prediction.** Three different scale windows are extracted, Small, Mid and Large, corresponding to $21\times21\,\mu m$, $42\times42\,\mu m$ and $84\times84\,\mu m$ FOV respectively. For the mid-scale, we also perform masking, by removing either the neighbor cells or the central cell to determine the important features for prediction. . . . . . . . . . . 3

E3 **Generative modeling of "synthetic" trajectories.** For each synthetic trajectory we start by encoding a real image as a starting point. Next, we take a random walk in the latent space. These trajectories in latent space are used as inputs to the TCN. Here, we also use the decoder to generate image sequences that represent the random walks in latent space. . . . . . . . . . 4

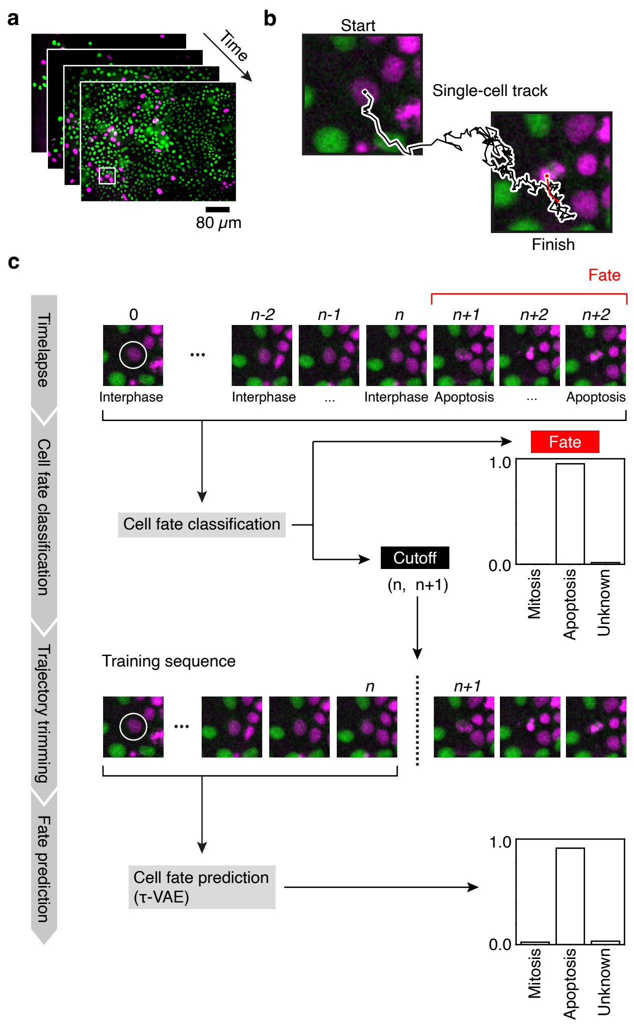


Figure E1: **Data flow in the model.** (a) Example time-lapse microscopy data showing a mixed population of MDCK$^{WT}$ (green) and scrib$^{kd}$ (magenta) cells. (b) Single-cell tracking is used to build a detailed training dataset of trajectories. The single-cell track is used to extract a glimpse of the cell over time, that becomes the input data for the machine learning models. (c) The data preparation and inference pipeline. A CNN/LSTM network classifies the fate of the cell and determines the cutoff point to truncate the track to remove images that encode the fate of the cell. The goal of the machine learning model is then to learn a representation that can predict the fate of a cell (circled in white) given the local configuration during interphase. Importantly, the model does not actually *observe* the fate, since these data fall beyond the cutoff. Images are taken at 4 minute intervals, MDCK$^{WT}$ cells appear in green and scrib$^{kd}$ in magenta

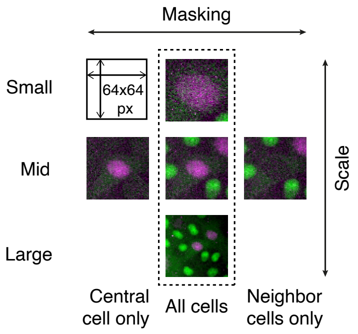


Figure E2: **Glimpse extraction and cell masking to determine the best image input scale for prediction.** Three different scale windows are extracted, Small, Mid and Large, corresponding to $21 \times 21\,\mu m$, $42 \times 42\,\mu m$ and $84 \times 84\,\mu m$ FOV respectively. For the mid-scale, we also perform masking, by removing either the neighbor cells or the central cell to determine the important features for prediction.

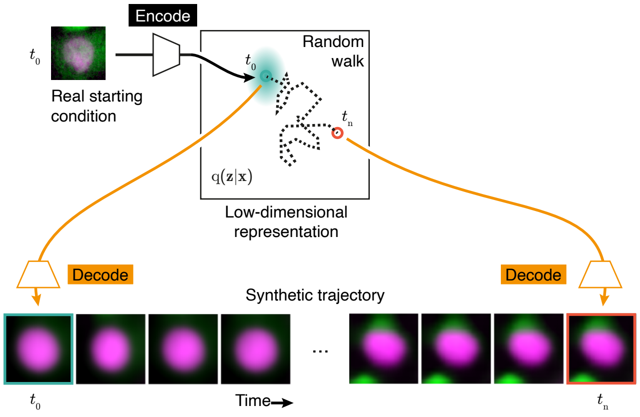


Figure E3: **Generative modeling of "synthetic" trajectories.** For each synthetic trajectory we start by encoding a real image as a starting point. Next, we take a random walk in the latent space. These trajectories in latent space are used as inputs to the TCN. Here, we also use the decoder to generate image sequences that represent the random walks in latent space.
