## Supplemental Data 10 for "Learning the Rules of Cell Competition without Prior Scientific Knowledge"

**Alan R. Lowe**[1,2,4,5*]

[1]Institute for the Physics of Living Systems
[2]Department of Structural and Molecular Biology
[3]Department of Cell and Developmental Biology
[4]London Centre for Nanotechnology
University College London, Gower St, London, WC1E 6BT, UK

[5]The Alan Turing Institute
Euston Rd, London NW1 2DB, UK

February 23, 2022

**2 Supplementary figures** **9**

**3 Supplementary tables** **28**

**4 Supplementary movies** **29**

S4 **Cumulative variance of principal components, in order of their explained variance ratio.** Principal component analysis was conducted on the latent spaces of four $\beta$-VAE models, with 8, 16, 32 and 64 latent dimensions respectively. The cumulative variance of the principal components was then plotted for each $\beta$-VAE. For a $\beta$-VAE with a latent dimension of $n$, we extracted $n$ principal components via PCA. Therefore, in the case of the "8dim" $\beta$-VAE, all of the variance is accounted for by 8 principal components, and so on. The principal components associated with the 8dim model reach a total variance of 65.83. For the 16dim, 32dim and 64dim models, this value is 68.33, 70.17 and 69.92 respectively. . 12

S5 **Example latent space traversals using the latent variables of the trained 32-dimensional $\beta$-VAE before PCA was applied.** Three latent variable pairs are represented: $z0$ with $z1$, $z1$ with $z2$ and $z2$ with $z3$. . . . . . . . 13

S6 **Graded examples of PC0 and PC1.** This figure displays some example cell fluorescence images that correspond to certain values of PC0 and PC1. For each component value $V = -4.0, 3.0, ..., 4.0$, twenty images are shown for whom their associated PC0 or PC1 value $v$ falls within the range $V - 0.5 < v < V + 0.5$. The sub-figure below shows the resulting "average image" obtained by taking the mean of every image in a 100,000-image dataset that corresponds to the aforementioned PC-value ranges. The number above each average image represents the number of raw images that have been averaged to obtain it. . . . . . . . 14

S7 **Examples of all Principal Components.** After PCA was applied to the latent space, it was found that several of the principal components corresponded to interpretable features of the cell fluorescence images. This figure portrays the result of taking the mean of all images from a dataset of 100,000 images whose corresponding value of a particular principal component fell within a specific range (the $x$-axis value ± 0.5). What is obtained is a way of visualising how the images differ with the variation of any one principal component. This visualisation is shown for all thirty-two principal components extracted using PCA. . . . . . . . 15

S8 **Correlation of all Principal Components with measureable parameters.** This figure shows the correlation coefficients of all the Principal Components with certain calculated variables. These variables were calculated based on the intensity images of example cell crops and their associated U-Net segmentations (see Section 1.9 for more details). The results shown here can be cross-referenced with the results shown in Figure S7 to arrive at an interpretation of the physical features to which the Principal Components correspond. . . . . . . . 16

S9 **Example scrib$^{\mathbf{kd}}$trajectories.** Each trajectory is cropped to roughly 30 minutes around the cutoff (red vertical line). The cutoff represents the point after which there are visible changes to the chromatin morphology which signify either apoptosis or mitosis. In some cases, the cutoff is placed several time-points before morphological change becomes visible (up to 30 minutes before). . . . . . . . 17

S10 **Generative modeling of “synthetic” trajectories.** For each synthetic trajectory we start by encoding a real image as a starting point. Next, we take a random walk in the latent space. These trajectories in latent space are used as inputs to the TCN. Here, we also use the decoder to generate image sequences that represent the random walks in latent space. Five example synthetic trajectories are shown. . . . . 18

S11 **Normalized confusion matrix for the testing results of using an LSTM backbone for cell fate prediction.** In order to establish a standard against which to assess the predictive performance of the TCN, we trained an LSTM that contained a very similar number of trainable parameters as the TCN (Section 1.11). 10-fold cross-validation was performed in order to obtain a result that was independent of the particular testing set chosen. The confusion matrix shown here is the result of averaging the testing results of LSTMs trained on the $\mathrm{MDCK}^{\mathrm{WT}}$and $\mathrm{scrib}^{\mathrm{kd}}$datasets respectively. . . . . . . . . . . 19

S12 **Normalized confusion matrices obtained when the $\tau$-VAE prediction model is tested on $\mathrm{MDCK}^{\mathrm{WT}}$trajectories from the competition data and uninduced data, respectively.** This figure allows comparison of the performance of the $\tau$-VAE on $\mathrm{MDCK}^{\mathrm{WT}}$trajectories from the different regimes. . . . . . . . . . . . . . . . . . . . . . . . . . . . . . . . . . . . . . . . . . . . . . . 20

S13 **Example of a feature saliency heatmaps for a $\mathrm{scrib}^{\mathrm{kd}}$ apoptosis event.** Here we calculate the feature saliency w.r.t. the input pixel data by backpropagating through the TCN and the convolutional encoder of the $\beta$-VAE. The input image data is shown in the left column. The middle column shows pixel saliency in the GFP channel of the input. The right column shows pixel saliency in the RFP channel of the input. Each image of the saliency is normalized per time point. Large gradient magnitudes (reds, yellows) indicate higher feature saliency. White arrows at indicate examples of regions of high saliency corresponding to nearby dividing cells or changes in the nuclear morphology of the target cell. . . . . 21

S14 **Fraction of correct predictions at different timescales for $\mathrm{scrib}^{\mathrm{kd}}$ cells.** After being fed a given number of frames of input, the TCN assigns a final-layer logit value to each label (apoptosis, mitosis or synthetic), which, after application of the softmax activation function, can be taken as the "confidence" value of the TCN in any particular label. In general, as the TCN is fed consecutive frames, it becomes gradually more confident in the correct prediction. This figure shows how the fraction of trajectories classified correctly with a threshold of $T = 0.90$ increases as the length of input increases. In other words, these plots show the fraction of trajectories for which the network predicts the correct fate with a confidence of at least 90%, for a given length of input. . . . . . . . . . . . . . . . . . . . . . . . . . 22

S15 **PC1 trajectories.** This figure shows the result of collecting all of the $\mathrm{scrib}^{\mathrm{kd}}$cell trajectories that were labelled 'apoptosis' or 'mitosis' (by the trajectory-classification network, in the case of the mitoses, and manually, in the case of the apoptoses) and then finding the average value over time of PC1, which correlates strongly with cell size/cell density. This was done for the "Competition", "BIRB" and "Uninduced" conditions. . . . . . . . . . . . . . . . . . . . . . . . . . . . . . . . . . . . . . . . . . 23

S16 **Normalized cell counts for $\mathrm{MDCK}^{\mathrm{WT}}$ and $\mathrm{scrib}^{\mathrm{kd}}$ cells under various conditions.** "Competition" (cell competition between $\mathrm{MDCK}^{\mathrm{WT}}$ and $\mathrm{scrib}^{\mathrm{kd}}$ cells), "BIRB" (competition in the presence of BIRB796), "Uninduced" (where the $\mathrm{scrib}^{\mathrm{kd}}$ cells are uninduced and therefore neither knock-down nor competition occur), and "DMSO" (competition in the presence of dimethyl sulfoxide). The ratio of $\mathrm{MDCK}^{\mathrm{WT}}$ to $\mathrm{scrib}^{\mathrm{kd}}$ cells at the beginning of the experiments was prepared to be 50:50. The cell counts are normalized relative to the initial count at the beginning of the experiment. This initial level is represented by the "Baseline" plot. . . . . . . . . . . . . . . . . . . . . . . . . . . . . . . . . . . 24

S17 **Example incorrect predictions of cell fate in the uninduced ($\mathrm{scrib}^{\mathrm{kd,\,tet-}}$) dataset.** Two example trajectories are shown, with one sub-figure for each. At the top of each sub-figure is placed a collage of time-points of the trajectory before the "cutoff" point. To the right is the final time-point of the trajectory (after the cutoff point), revealing the cell fate. Below that is shown, in order: the confidence plot, showing the TCN's predictions over time; a plot of PC1, the most important principal component for fate prediction; a saliency heat-map showing which components were most important to the prediction, and when and; a saliency plot over time for PC1. . . . . . . . . . . . . . . . . . . . . . . . . . 25

S18 **Example incorrect predictions of cell fate in the BIRB796 treated dataset.** Two example trajectories are shown, with one sub-figure for each. At the top of each sub-figure is placed a collage of time-points of the trajectory before the "cutoff" point. To the right is the final time-point of the trajectory (after the cutoff point), revealing the cell fate. Below that is shown, in order: the confidence plot, showing the TCN's predictions over time; a plot of PC1, the most important principal component for fate prediction; a saliency heat-map showing which components were most important to the prediction, and when and; a saliency plot over time for PC1. . . . . . . . . . . . . . . . . . . . . . . . . . . . . . . . . 26

S19 **Normalized confusion matrix for the testing of a neural network trained to discriminate between the final-frame encodings of mitotic, apoptotic and "other" trajectories.** This model was trained for 100 epochs, and its architecture consisted of three dense layers with 64, 128 and 256 units respectively, followed by a final logits layer with 3 units. . . . . . . . . . . . . . . . . . . . . . . . . . . 27

#### List of Tables

1 Datasets used in this study. All movies were captured with a size 1600×1200 pixels (corresponding to 530×400 µm) with 3 channels (Brightfield, GFP and RFP). . . . . . . . . . . . . . . . . . . . . . . . . 28
2 Single-cell trajectory training datasets used in this study. († see Methods) . . . . . . . . . . . . . . . . 28
3 Drugs used in this study . . . . . . . . . . . . . . . . . . . . . . . . . . . . . . . . . . . . . . . . . 28
4 The testing macro-F1 of trained TCNs using various VAE models. . . . . . . . . . . . . . . . . . . . . 28

### 1 Additional methods

#### 1.1 Cell culture

The MDCK cell lines used for this study scrib$^{kd}$ were a kind gift from Prof Yasuyuki Fujita (University of Kyoto, Japan) and described in [1].To enable visualisation of nucleic acid organisation during the cell cycle, we established cell lines stably expressing fluorescently tagged histone markers. Use of different fluorescent proteins enabled us to distinguish the two competing cell types and allowed for accurate segmentation. MDCK$^{WT}$ and scrib$^{kd}$ cells expressing H2B-GFP and -RFP nuclear markers were described in [Bove et al, 2017]. All cell lines used in this publication have been tested for mycoplasma infection and were found to be negative (MycoAlert Plus Detection Kit, Lonza, LT07-710). MDCK cells were grown in DMEM (Thermo-Fisher) supplemented with 10% fetal bovine serum (FBS, Sigma-Aldrich), Hepes buffer (Sigma-Aldrich), and 1% Penicillin/Streptomycin in a humidified incubator at 37°C with 5% $CO_2$. The scrib$^{kd}$ cells were cultured as wild-type cells, except that we used tetracycline-free bovine serum (Clontech, 631106) to supplement the culture medium. For inducing expression of scribble shRNA, doxycycline (Sigma-Aldrich, D9891-1G) was added to the medium at a final concentration of 1 $\mu$g/ml.

#### 1.2 Automated Widefield Microscopy

A custom-built automated epifluorescence microscope was built inside a standard $CO_2$ incubator (Heraeus BL20) which maintained the temperature at 37°C and 5% $CO_2$. The microscope utilised an 20× air objective (Olympus Plan Fluorite, 0.5 NA, 2.1mm WD), high performance encoded motorized XY and focus motor stages (Prior H117E2IX, FB203E and ProScan III controller) and a 9.1MP CCD camera (Point Grey GS3-U3-91S6M). Brightfield illumination was provided by a fibre-coupled green LED (Thorlabs, 530nm). GFP and mCherry/RFP fluorescence excitation was provided by a LED light engine (Bluebox Optics niji). Cameras and light sources were synchronised using TTL pulses from an external D/A converter (Data Translation DT9834). Sample humidity was maintained using a custom built chamber humidifier. The microscope was controlled using MICRO-MANAGER [2] and our own software OCTOPUSLITE[2].

#### 1.3 Cell competition assay

Cell competition assays were carried out in 24-well imaging plates (ibidi). At the start of each experiment, cells were seeded at an initial density of $1 \times 10^{-3}$ cells/$\mu m^2$. MDCK$^{WT}$ cells expressing H2B-GFP were mixed with scrib$^{kd}$ H2B-RFP cells at a ratio of 90:10, 50:50 or 10:90. In some experiments, the expression of scribble shRNA was been induced in scrib$^{kd}$ cells by exposure to 1ug/mL doxycycline for 70 hours before seeding. In other experiments, the cells were maintained in tetracycline free medium to prevent scribble shRNA induction. Imaging was started 2–3 h after seeding. Imaging medium used during the assay was phenol red free DMEM (Thermo Fisher Scientific, 31053) supplemented with tetracycline-free bovine serum, Hepes, antibiotics and, for experiments involving induction, doxycycline at the dose indicated above. Multi-location imaging was performed inside the incubator-scope acquiring Brightfield, GFP and RFP fluorescence images with a frequency of 1 frame every 4 minutes for each position for 80 hours (1200 frames).

#### 1.4 Image alignment and normalization

Image stacks were aligned using StackReg [3]. All images used in the analysis were normalized to have a mean value of zero and unit variance.

#### 1.5 Single cell tracking

Instance segmentation of individual cells in timelapse microscopy sequences was performed using a residual U-Net as previously described. We used a Bayesian cell tracking approach[3] to assemble single-cell cell trajectories from the data [4, 5].

#### 1.6 Glimpse extraction, cell fate classification and determination of cutoff

Glimpses are extracted at $64 \times 64$, $128 \times 128$ and $256 \times 256$ pixels, centred on the cell of interest at each time point, and downsampled to $64 \times 64$ pixels. We use the first scale as input to the trajectory classification network, reasoning that the morphology of the nucleus is sufficient to classify the fate of the cell. We trained a combined CNN-LSTM

[2]https://www.github.com/quantumjot/octopuslite
[3]https://www.github.com/quantumjot/bayesiantracker

neural network to classify each trajectory as containing either a mitotic or an apoptotic event. This network was trained using 7,922 manually labelled trajectories: 4,401 mitoses, 828 apoptoses and 2,693 "unknown" trajectories. The latter refers to trajectories in which neither a mitosis nor an apoptosis occurs. The CNN and the LSTM [6] were trained separately. The CNN was trained to classify individual fluorescence images according to the morphological state of the nucleus ("interphase", "metaphase", "prometaphase", "anaphase" or "apoptotic") [4, 5]. The LSTM was trained to integrate the time-specific state classifications in a trajectory to arrive at a classification of the trajectory as a whole ("mitotic", "apoptotic" or "unknown"). Trajectories terminating with an unknown classification arise when a cell exits the microscope's field of view (FOV) or the movie terminates before an event can occur.

Various augmentation techniques were used to train both the CNN and the LSTM. For the CNN, we used random flipping, cropping, rotation, translation, noise addition, brightness adjustment, and aspect ratio adjustment on the training images. For the LSTM, we used frame-swapping, random-frame-deletion and noise addition on the training trajectories. After all of the labelled trajectories were obtained, an automated procedure was applied to shorten the duration of the glimpses so as to leave out the portion of each time-series that contains the fate event (the mitosis or apoptosis event) (**Fig S8**). This pruning procedure ensured that the prediction network could not use morphological changes arising as a consequence of cell fate.

##### 1.7 Confusion matrices

To assess the classification performance of our networks, we computed a confusion matrix. In a confusion matrix, the number in row *i* and column *i* represents the number of data instances in the testing set that are of ground-truth class *i* yet are classified as belonging to class *j* by the model. We often used a normalized confusion matrix, where each element in row *i* is written as a proportion of the sum of elements in row *j* (i.e. the number of ground-truth examples of class *j*).

##### 1.8 Cell masking procedure

We performed cell masking as follows. First, we used the U-Net segmentation of our glimpse images to find the regions in each image that corresponded to cells, and conversely, the regions that corresponded to the background. Next, we identified the cell-regions that corresponded to either the central cell, or the cells in the neighbourhood, depending on which we wished to mask. In the "Mid-View, Central Cell Only" framework, we masked the neighbourhood cells, whereas in the "Mid-View, Neighbour Cells Only" framework we masked the central cell. Finally, we calculated the mean and standard deviation of pixel values in the background region, and then used these values to replace the masked region with Gaussian noise, such that this region appears similar to the background.

##### 1.9 Calculation of image properties and correlation with principal components

To calculate image properties, we used SCIKIT-IMAGE regionprops. We calculated the following properties for the central cell of each glimpse using the intensity images and the U-Net segmentation:

- `area`
- `eccentricity`
- `orientation`
- `solidity`
- `intensity_mean`

Separately, we also calculated the number of cells by counting the number of unique connected components in the U-Net binary segmentation of the glimpse, using the `label` function from the `morphology` module of scikit-image. The aspect ratio of the cells was calculated by finding the ratio between the maximum horizontal and vertical spans of the cells as given by their U-Net segmentations.

##### 1.10 Covariance matrices

A $d \times d$ covariance matrix displays the joint variability of two variables $z_i$ and $z_j$ for $i, j \in \mathbb{N}, 0 \leq i, j \leq d$. The covariance of variables $z_i$ and $z_j$ is shown in row $i$ and column $j$ of the matrix. The covariance of $z_i$ and $z_j$ across a sample of size $n$ is defined by:

$$cov(z_i, z_j) = \frac{\sum_i^n (x_i - \overline{x}) \times (y_i - \overline{y})}{n - 1} \quad (1)$$

##### 1.11 Alternative $\tau$-VAE using a Long-Short Term Memory (LSTM) backbone

The LSTM is a form of Recurrent Neural Network (RNN) proposed by Hochreiter & Schmidhuber [6]. Its wide usage across many years made it the ideal benchmark against which to compare our $\tau$-VAE model. In essence, the LSTM works by processing each time-step of the input sequentially, as with any type of RNN. The difference between an LSTM and a "vanilla" RNN is that the LSTM maintains a "cell state" that interacts with the sequential inputs via a set of gates. These gates control how the inputs affect the cell state, and vice versa. In this way, the LSTM controls the information that passes on from time-step $t$ to time-step $t+1$. It is this control that enables the LSTM to integrate information over a large number of time-steps, a task in which vanilla RNNs face considerable difficulty.

The LSTM model we built was composed of three LSTM layers – with 32, 64 and 128 units respectively – followed by one dense layer with 32 units, and then a final logits layer with 3 units, one for each class. This model was trained with a batch size of 128 for 100 epochs using the RMSprop optimizer with a learning rate of 0.001 (the same procedure as the $\tau$-VAE model).

##### 1.12 Computational hardware

All code was implemented in Python and C/C++ using CVXOPT, GLPK, Numpy, Scipy, Scikit-Image, Scikit-Learn, TensorFlow, Keras and JAX libraries. Microscopy image visualisation was performed using NAPARI[7]. All image processing was performed on a ASUS ESC4000 G3 server (RackServers.com) running Ubuntu 18.04 LTS with 256Gb RAM and NVIDIA GTX1080 Ti or V100 GPUs.

#### 2 Supplementary figures

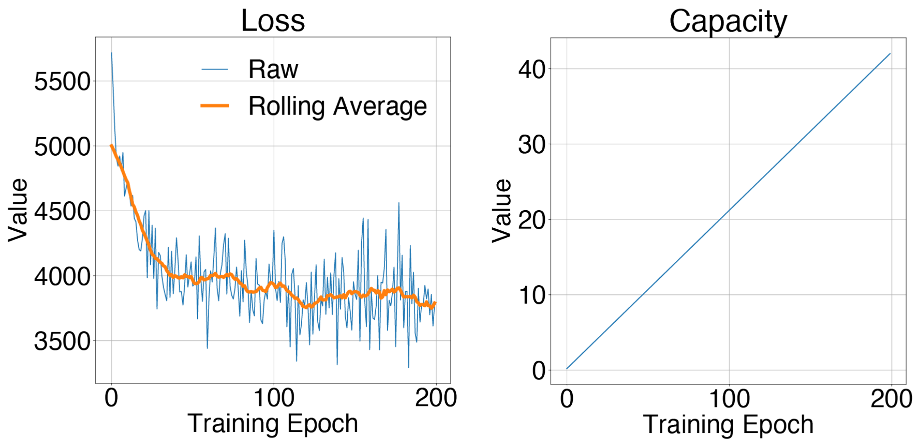

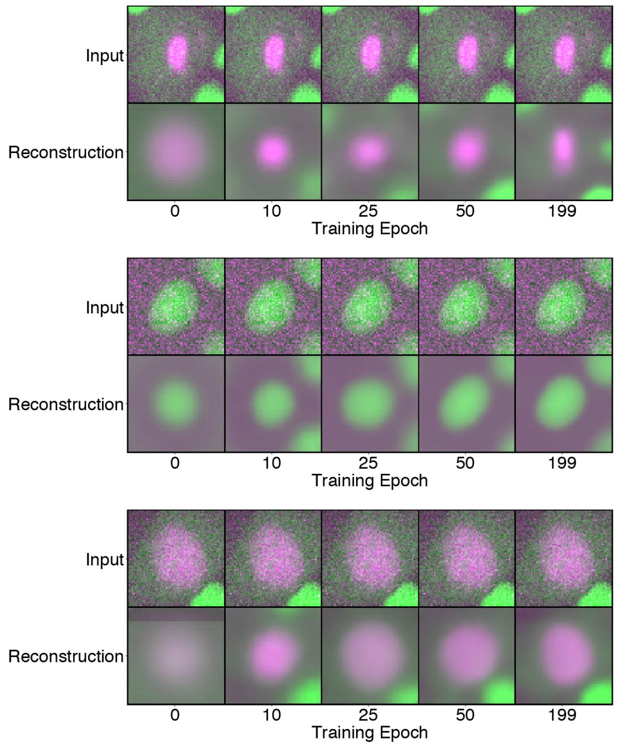

Figure S1: **Example of linear increase of $\beta$-VAE bottleneck capacity during training.** (a) Decreasing loss as a function of training iteration. (b) Linear increase in bottleneck capacity during training (d) Example images generated by sampling during training, demonstrating the acquisition of fine details as capacity increases.

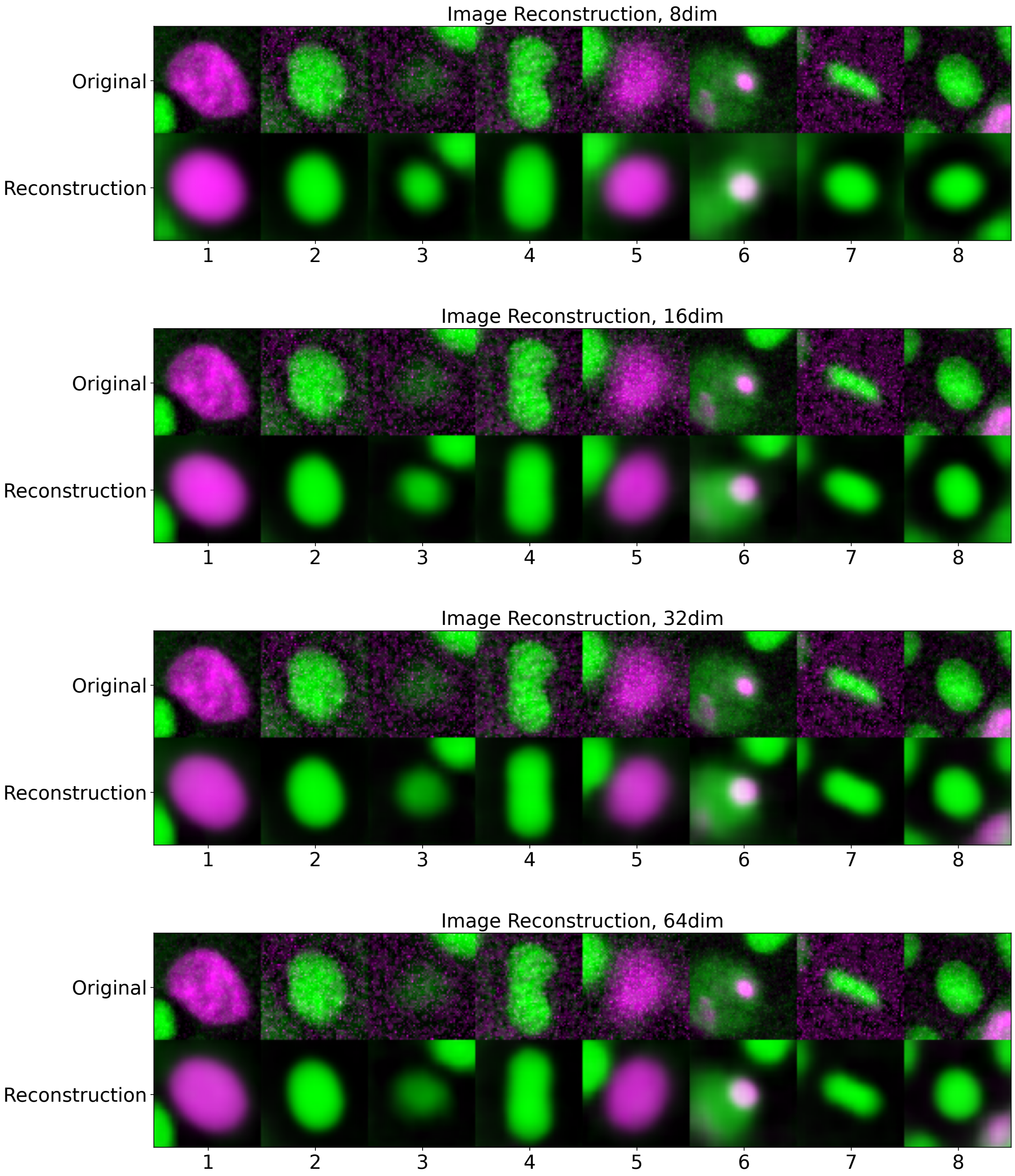

Figure S2: **Example image reconstructions using models of $\beta$-VAE that differ only in the size of their latent space.** Eight images were randomly chosen from the image dataset and then reconstructed using the four models.

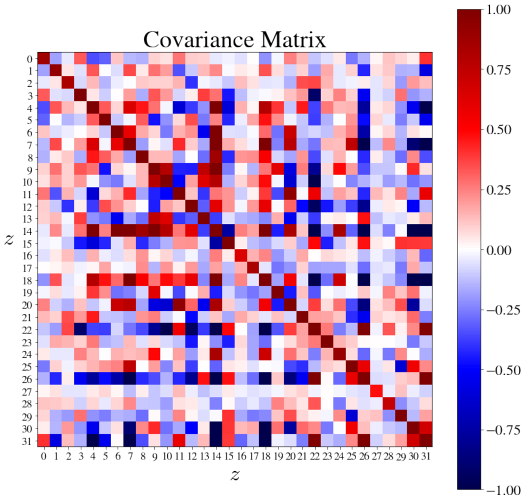

(a) **Raw latent variables.**

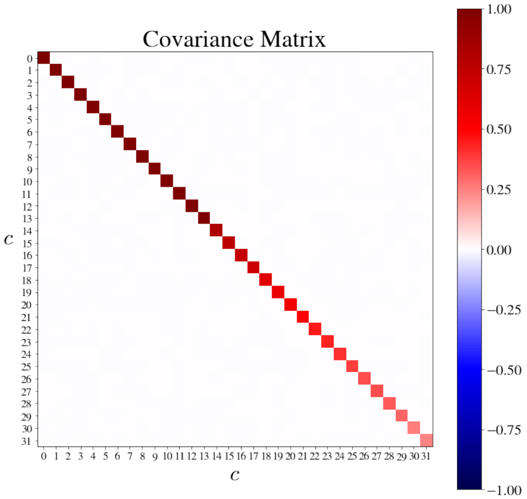

(b) **Principal components of latent variables.**

Figure S3: **Covariance matrices for the latent variables both pre- and post-PCA.** (a) A covariance matrix was plotted for the variables that form the latent space of the $\beta$-VAE. This matrix represents the degree to which two latent variables co-vary across a dataset of 1.2 million images taken from the cell culture (see 1.10 for a description of how covariance was calculated). (b) A covariance matrix was was then plotted for the principal components of the latent variables.

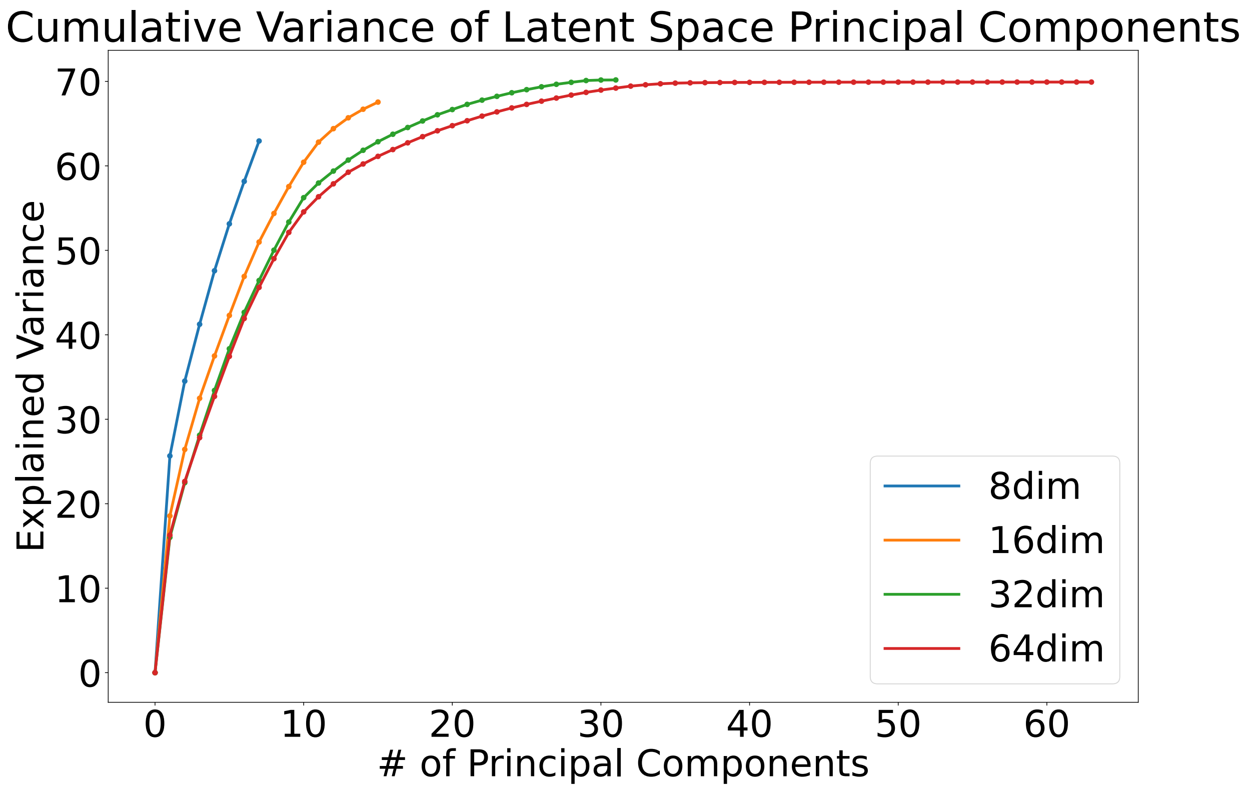

Figure S4: **Cumulative variance of principal components, in order of their explained variance ratio.** Principal component analysis was conducted on the latent spaces of four $\beta$-VAE models, with 8, 16, 32 and 64 latent dimensions respectively. The cumulative variance of the principal components was then plotted for each $\beta$-VAE. For a $\beta$-VAE with a latent dimension of $n$, we extracted $n$ principal components via PCA. Therefore, in the case of the "8dim" $\beta$-VAE, all of the variance is accounted for by 8 principal components, and so on. The principal components associated with the 8dim model reach a total variance of 65.83. For the 16dim, 32dim and 64dim models, this value is 68.33, 70.17 and 69.92 respectively.

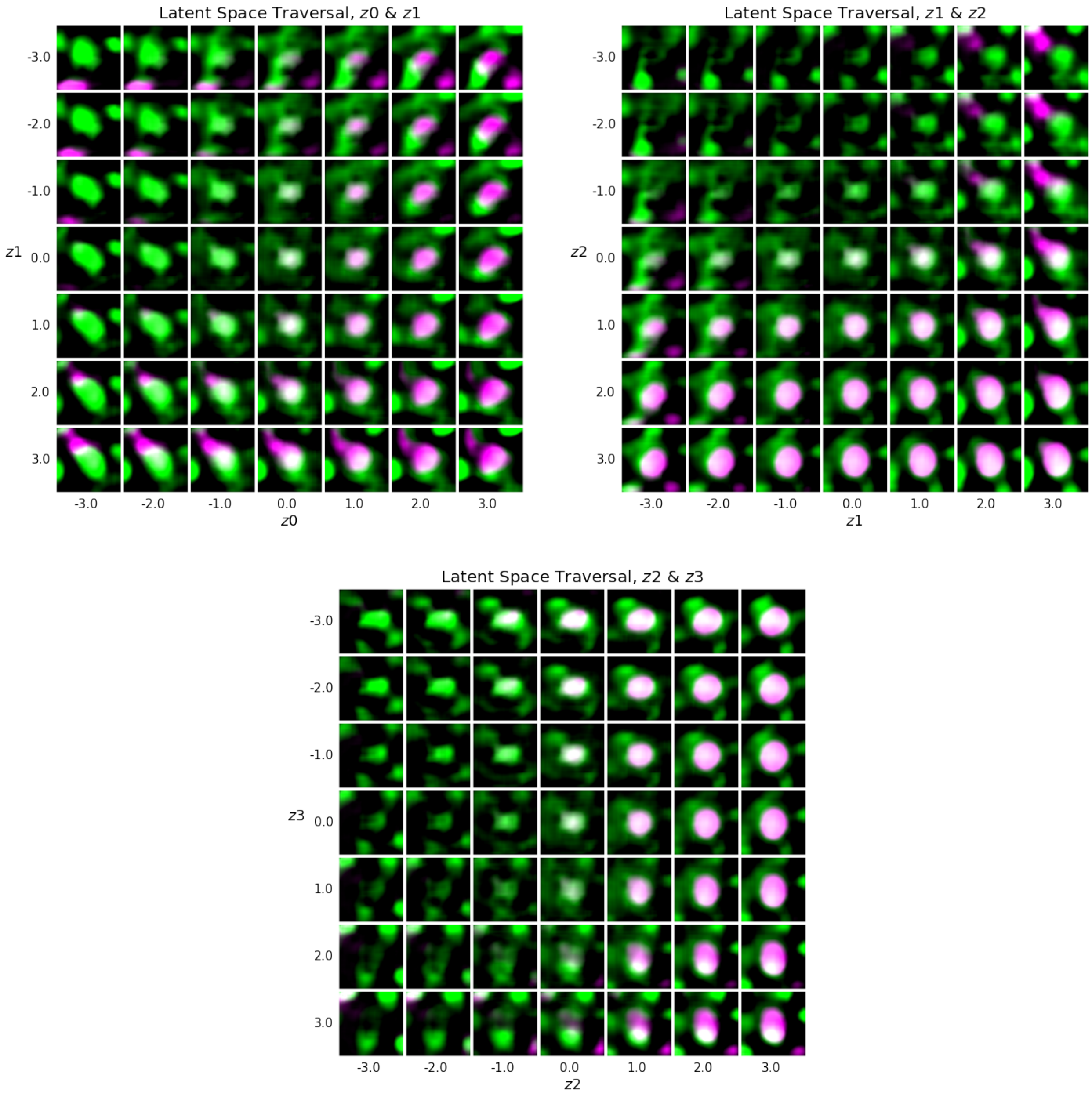

Figure S5: **Example latent space traversals using the latent variables of the trained 32-dimensional $\beta$-VAE before PCA was applied.** Three latent variable pairs are represented: $z0$ with $z1$, $z1$ with $z2$ and $z2$ with $z3$.

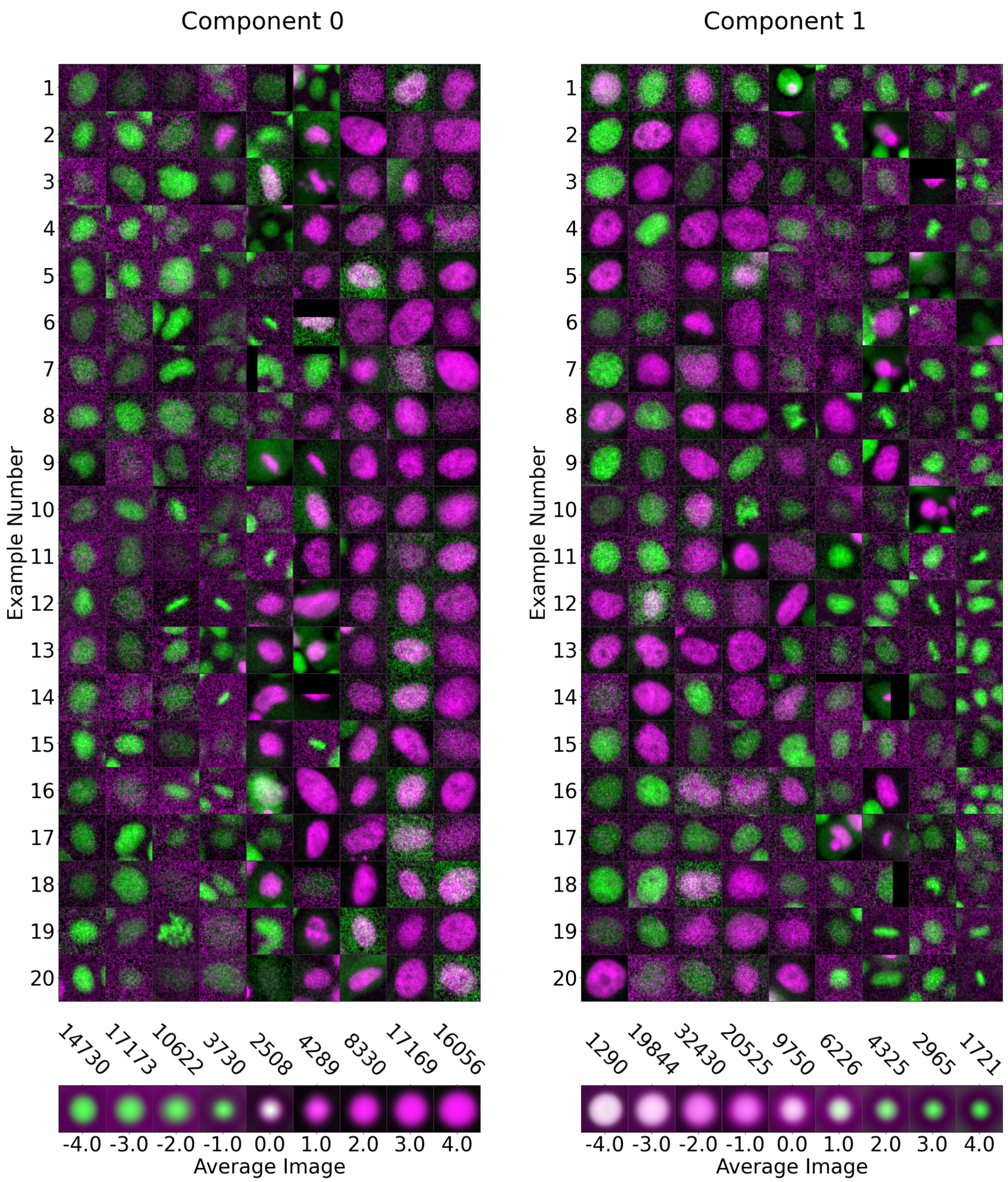

Figure S6: **Graded examples of PC0 and PC1.** This figure displays some example cell fluorescence images that correspond to certain values of PC0 and PC1. For each component value $V = -4.0, 3.0, ..., 4.0$, twenty images are shown for whom their associated PC0 or PC1 value $v$ falls within the range $V - 0.5 < v < V + 0.5$. The sub-figure below shows the resulting "average image" obtained by taking the mean of every image in a 100,000-image dataset that corresponds to the aforementioned PC-value ranges. The number above each average image represents the number of raw images that have been averaged to obtain it.

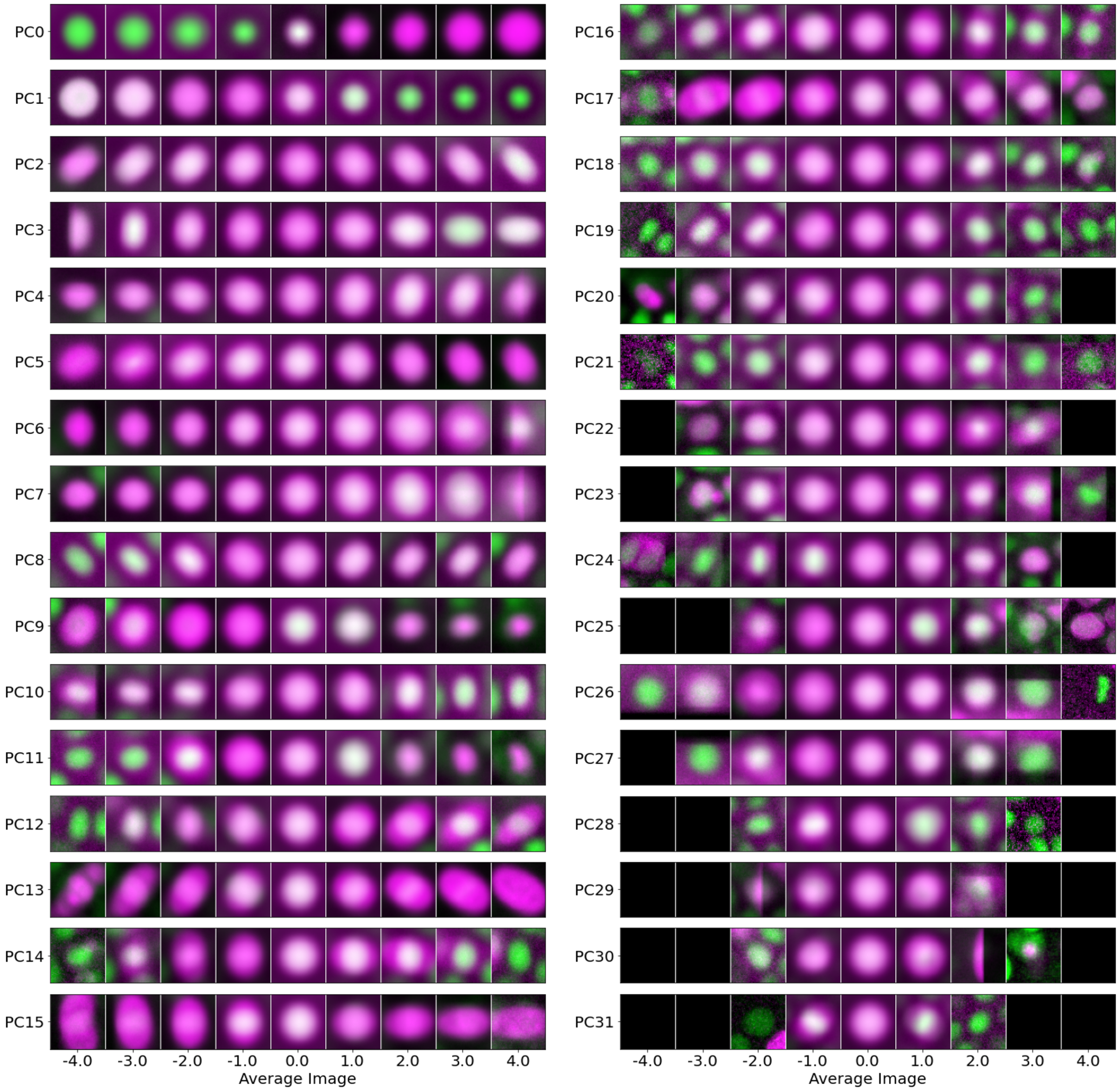

Figure S7: **Examples of all Principal Components.** After PCA was applied to the latent space, it was found that several of the principal components corresponded to interpretable features of the cell fluorescence images. This figure portrays the result of taking the mean of all images from a dataset of 100,000 images whose corresponding value of a particular principal component fell within a specific range (the $x$-axis value ± 0.5). What is obtained is a way of visualising how the images differ with the variation of any one principal component. This visualisation is shown for all thirty-two principal components extracted using PCA.

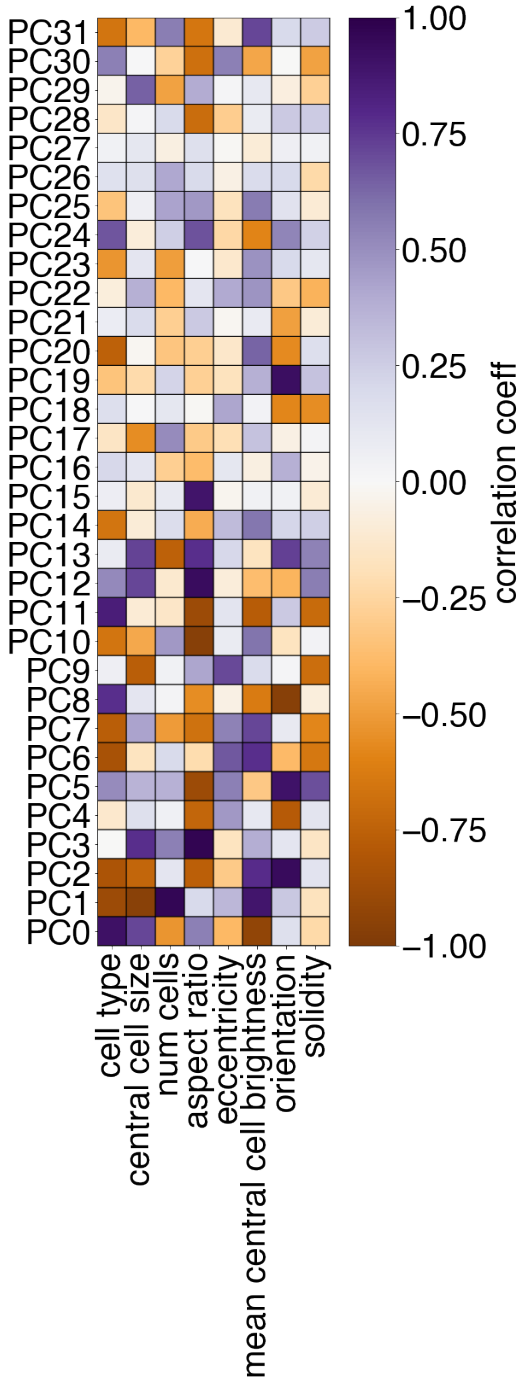

Figure S8: **Correlation of all Principal Components with measureable parameters.** This figure shows the correlation coefficients of all the Principal Components with certain calculated variables. These variables were calculated based on the intensity images of example cell crops and their associated U-Net segmentations (see Section 1.9 for more details). The results shown here can be cross-referenced with the results shown in Figure S7 to arrive at an interpretation of the physical features to which the Principal Components correspond.

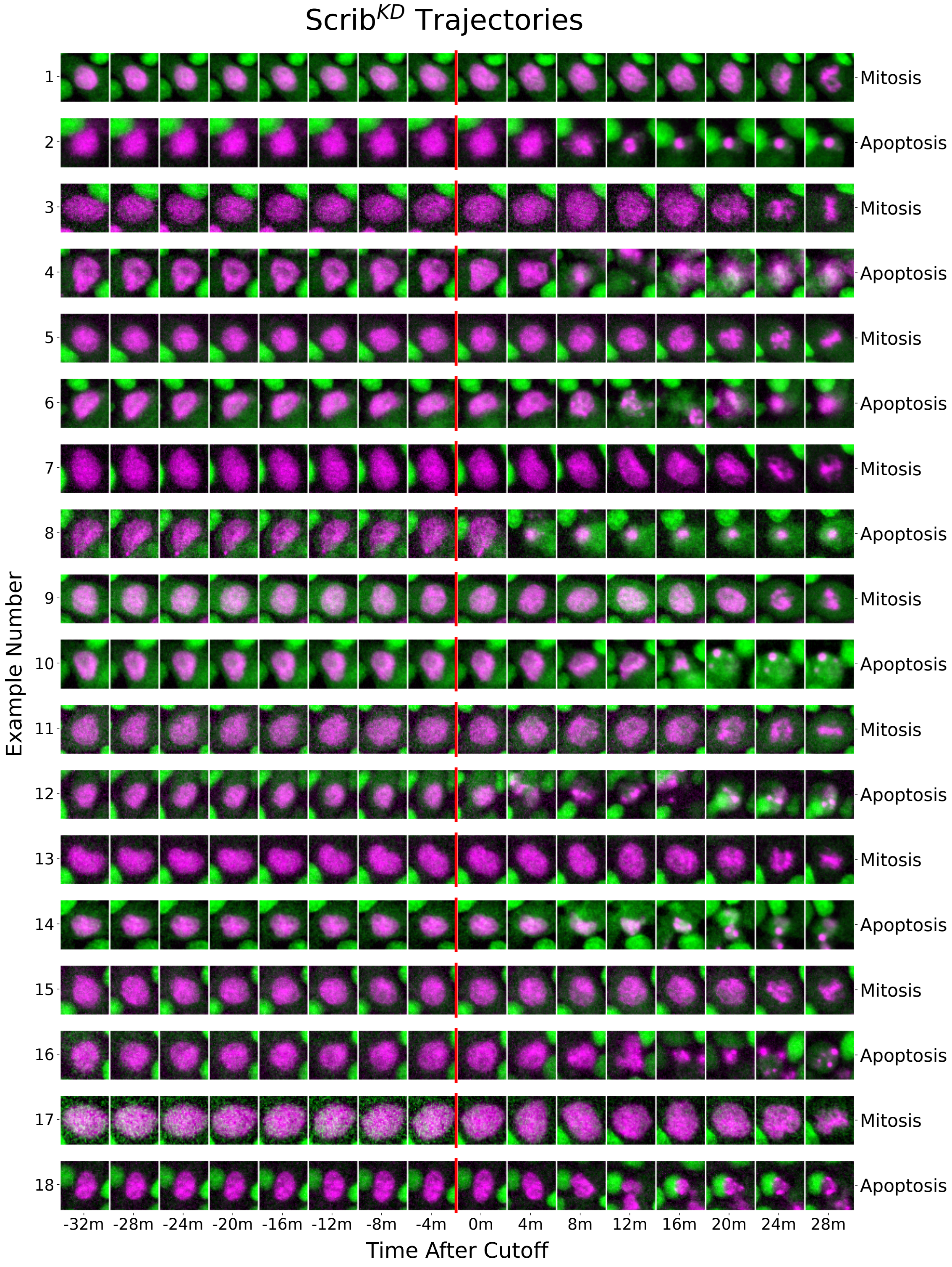

Figure S9: **Example scrib$^{kd}$ trajectories.** Each trajectory is cropped to roughly 30 minutes around the cutoff (red vertical line). The cutoff represents the point after which there are visible changes to the chromatin morphology which signify either apoptosis or mitosis. In some cases, the cutoff is placed several time-points before morphological change becomes visible (up to 30 minutes before).

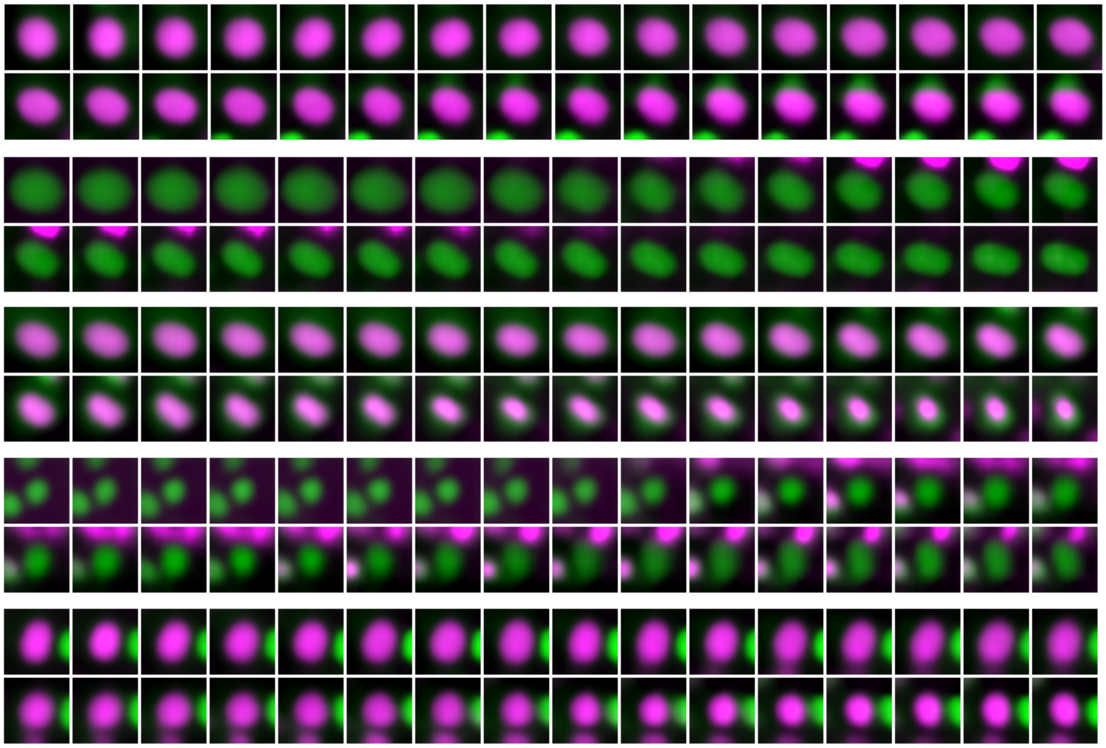

Figure S10: **Generative modeling of "synthetic" trajectories.** For each synthetic trajectory we start by encoding a real image as a starting point. Next, we take a random walk in the latent space. These trajectories in latent space are used as inputs to the TCN. Here, we also use the decoder to generate image sequences that represent the random walks in latent space. Five example synthetic trajectories are shown.

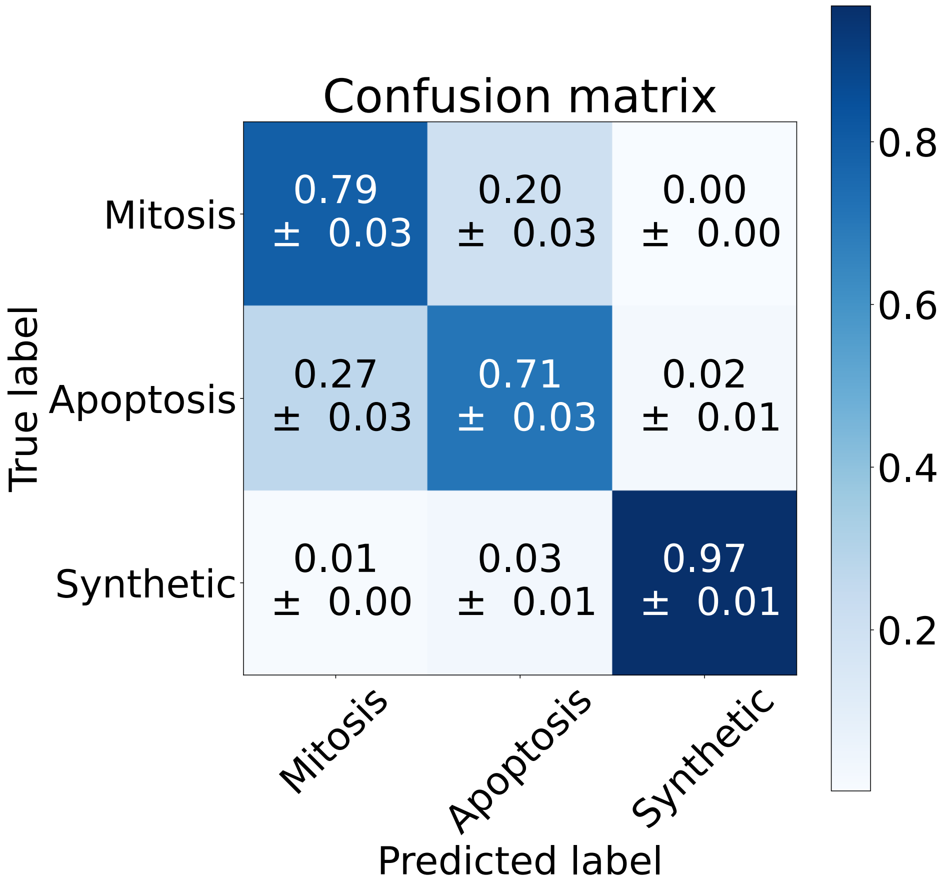

Figure S11: **Normalized confusion matrix for the testing results of using an LSTM backbone for cell fate prediction.** In order to establish a standard against which to assess the predictive performance of the TCN, we trained an LSTM that contained a very similar number of trainable parameters as the TCN (Section 1.11). 10-fold cross-validation was performed in order to obtain a result that was independent of the particular testing set chosen. The confusion matrix shown here is the result of averaging the testing results of LSTMs trained on the MDCK$^{WT}$and scrib$^{kd}$datasets respectively.

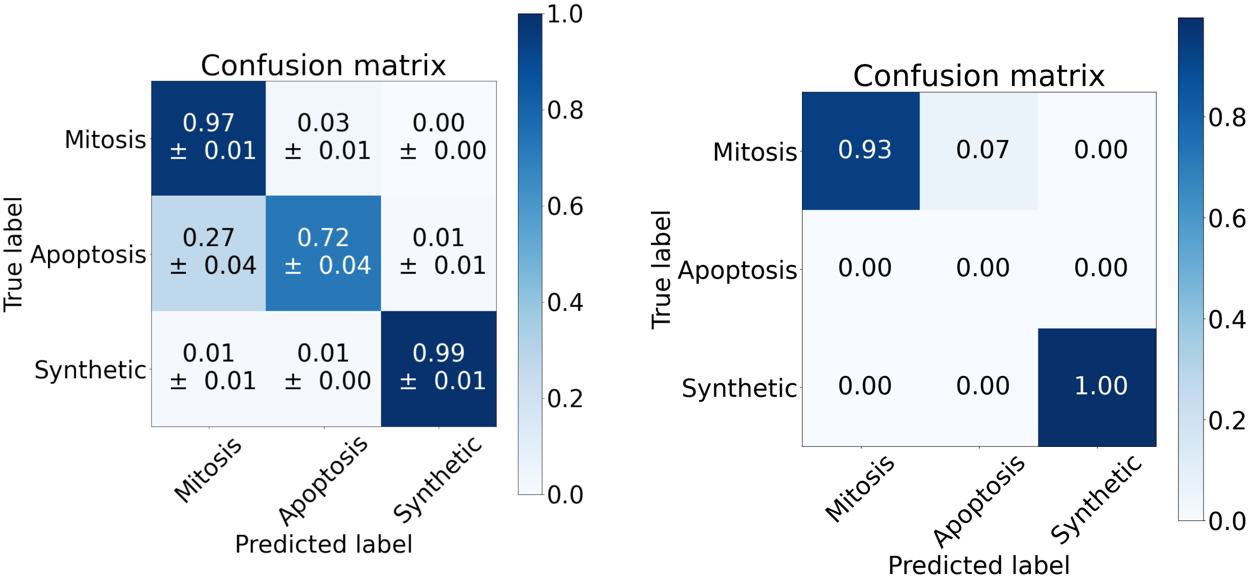

(a) Competition (MDCK$^{\mathrm{WT}}$:scrib$^{\mathrm{kd}}$).

(b) Uninduced (MDCK$^{\mathrm{WT}}$:scrib$^{\mathrm{kd,\ tet-}}$).

Figure S12: **Normalized confusion matrices obtained when the $\tau$-VAE prediction model is tested on MDCK$^{\mathrm{WT}}$trajectories from the competition data and uninduced data, respectively.** This figure allows comparison of the performance of the $\tau$-VAE on MDCK$^{\mathrm{WT}}$trajectories from the different regimes.

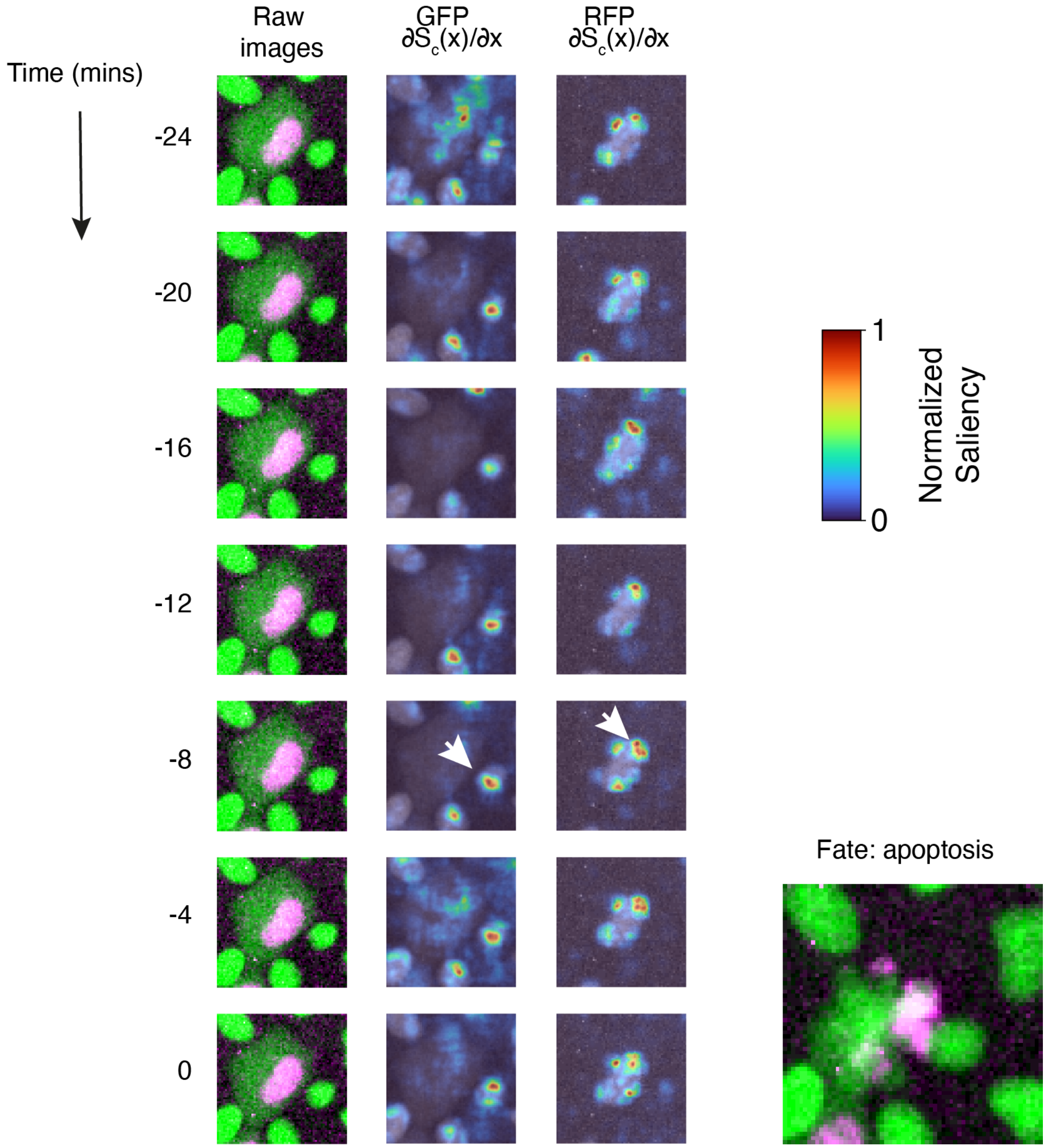

Figure S13: **Example of a feature saliency heatmaps for a scrib$^{kd}$ apoptosis event.** Here we calculate the feature saliency w.r.t. the input pixel data by backpropagating through the TCN and the convolutional encoder of the $\beta$-VAE. The input image data is shown in the left column. The middle column shows pixel saliency in the GFP channel of the input. The right column shows pixel saliency in the RFP channel of the input. Each image of the saliency is normalized per time point. Large gradient magnitudes (reds, yellows) indicate higher feature saliency. White arrows at indicate examples of regions of high saliency corresponding to nearby dividing cells or changes in the nuclear morphology of the target cell.

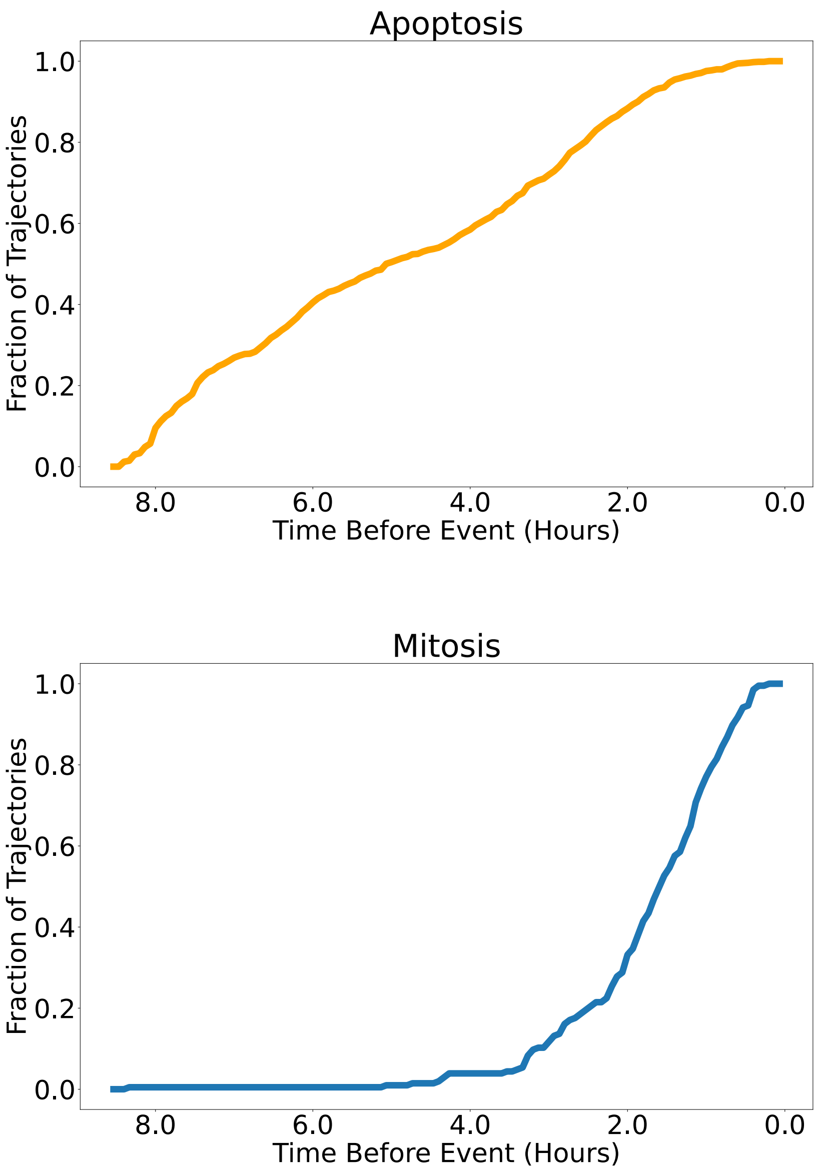

Figure S14: **Fraction of correct predictions at different timescales for scrib$^{kd}$ cells.** After being fed a given number of frames of input, the TCN assigns a final-layer logit value to each label (apoptosis, mitosis or synthetic), which, after application of the softmax activation function, can be taken as the "confidence" value of the TCN in any particular label. In general, as the TCN is fed consecutive frames, it becomes gradually more confident in the correct prediction. This figure shows how the fraction of trajectories classified correctly with a threshold of $T = 0.90$ increases as the length of input increases. In other words, these plots show the fraction of trajectories for which the network predicts the correct fate with a confidence of at least 90%, for a given length of input.

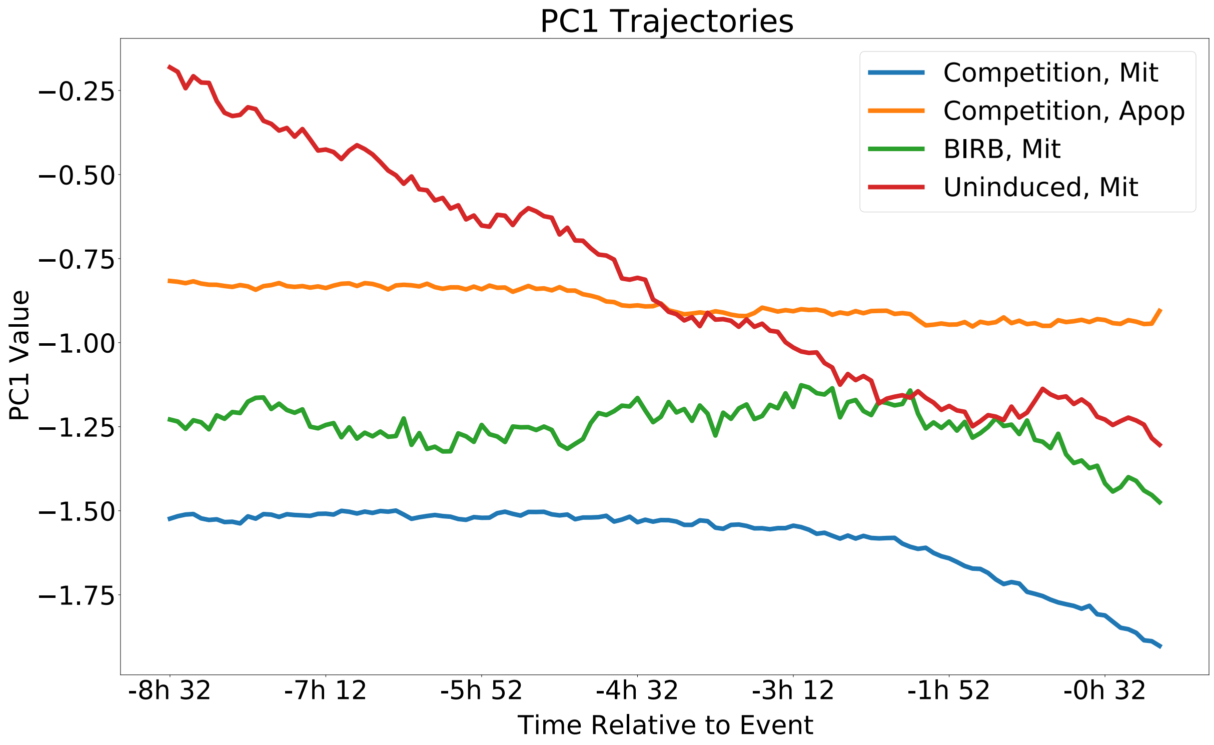

Figure S15: **PC1 trajectories.** This figure shows the result of collecting all of the scrib$^{kd}$cell trajectories that were labelled 'apoptosis' or 'mitosis' (by the trajectory-classification network, in the case of the mitoses, and manually, in the case of the apoptoses) and then finding the average value over time of PC1, which correlates strongly with cell size/cell density. This was done for the "Competition", "BIRB" and "Uninduced" conditions.

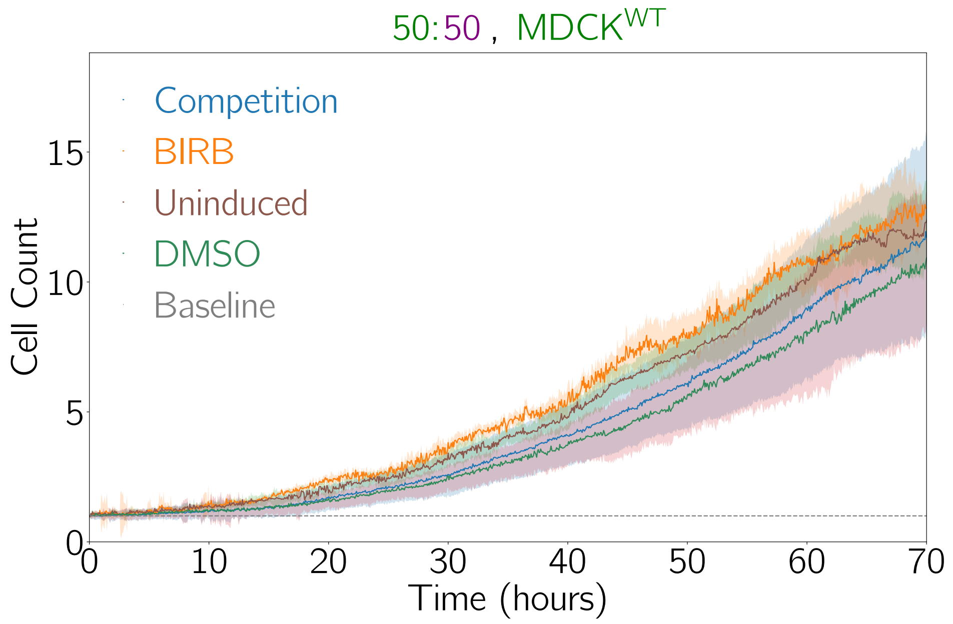

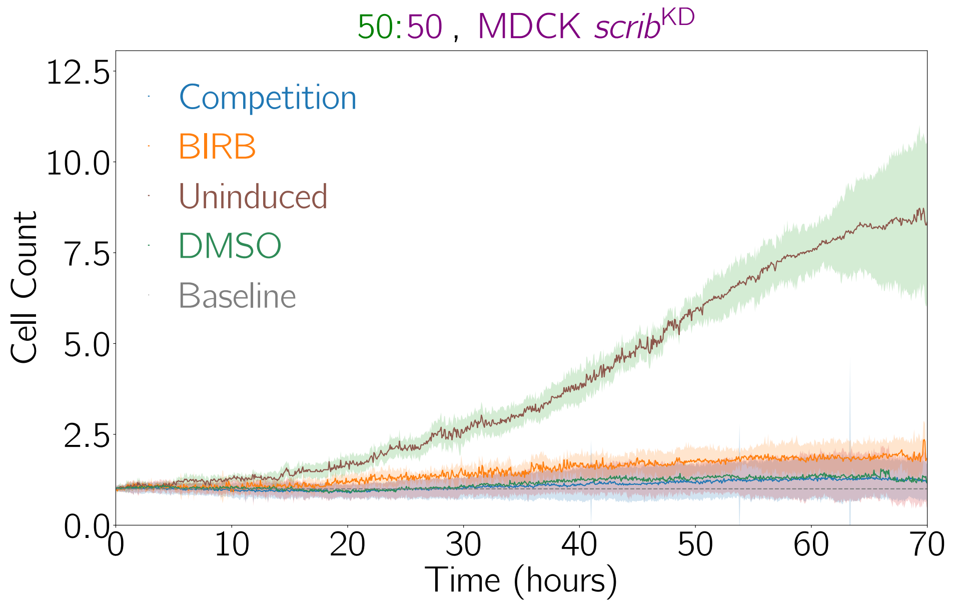

Figure S16: **Normalized cell counts for MDCK$^{WT}$ and scrib$^{kd}$ cells under various conditions.** "Competition" (cell competition between MDCK$^{WT}$ and scrib$^{kd}$ cells), "BIRB" (competition in the presence of BIRB796), "Uninduced" (where the scrib$^{kd}$ cells are uninduced and therefore neither knock-down nor competition occur), and "DMSO" (competition in the presence of dimethyl sulfoxide). The ratio of MDCK$^{WT}$ to scrib$^{kd}$ cells at the beginning of the experiments was prepared to be 50:50. The cell counts are normalized relative to the initial count at the beginning of the experiment. This initial level is represented by the "Baseline" plot.

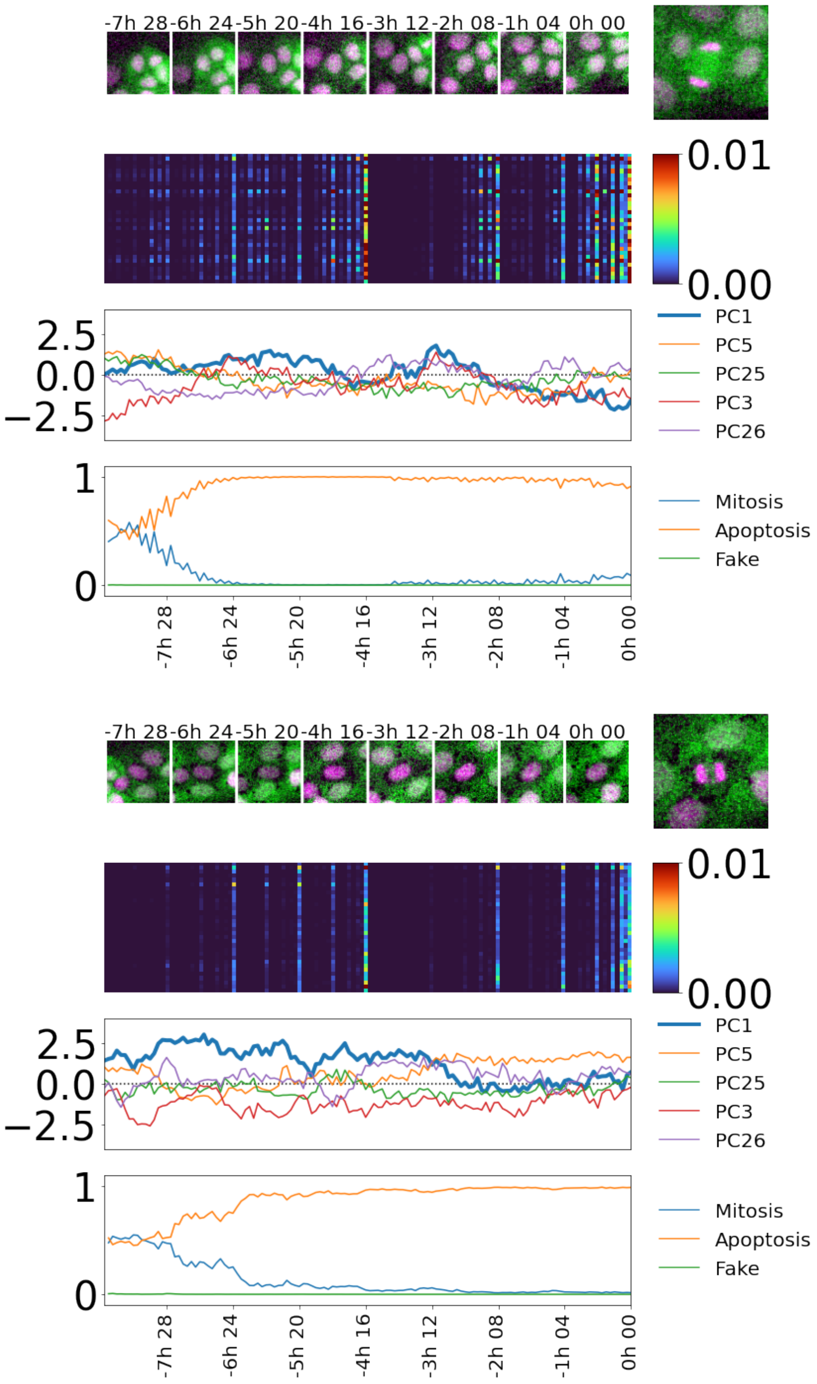

Figure S17: **Example incorrect predictions of cell fate in the uninduced (scrib$^{kd, tet-}$) dataset.** Two example trajectories are shown, with one sub-figure for each. At the top of each sub-figure is placed a collage of time-points of the trajectory before the "cutoff" point. To the right is the final time-point of the trajectory (after the cutoff point), revealing the cell fate. Below that is shown, in order: the confidence plot, showing the TCN's predictions over time; a plot of PC1, the most important principal component for fate prediction; a saliency heat-map showing which components were most important to the prediction, and when and; a saliency plot over time for PC1.

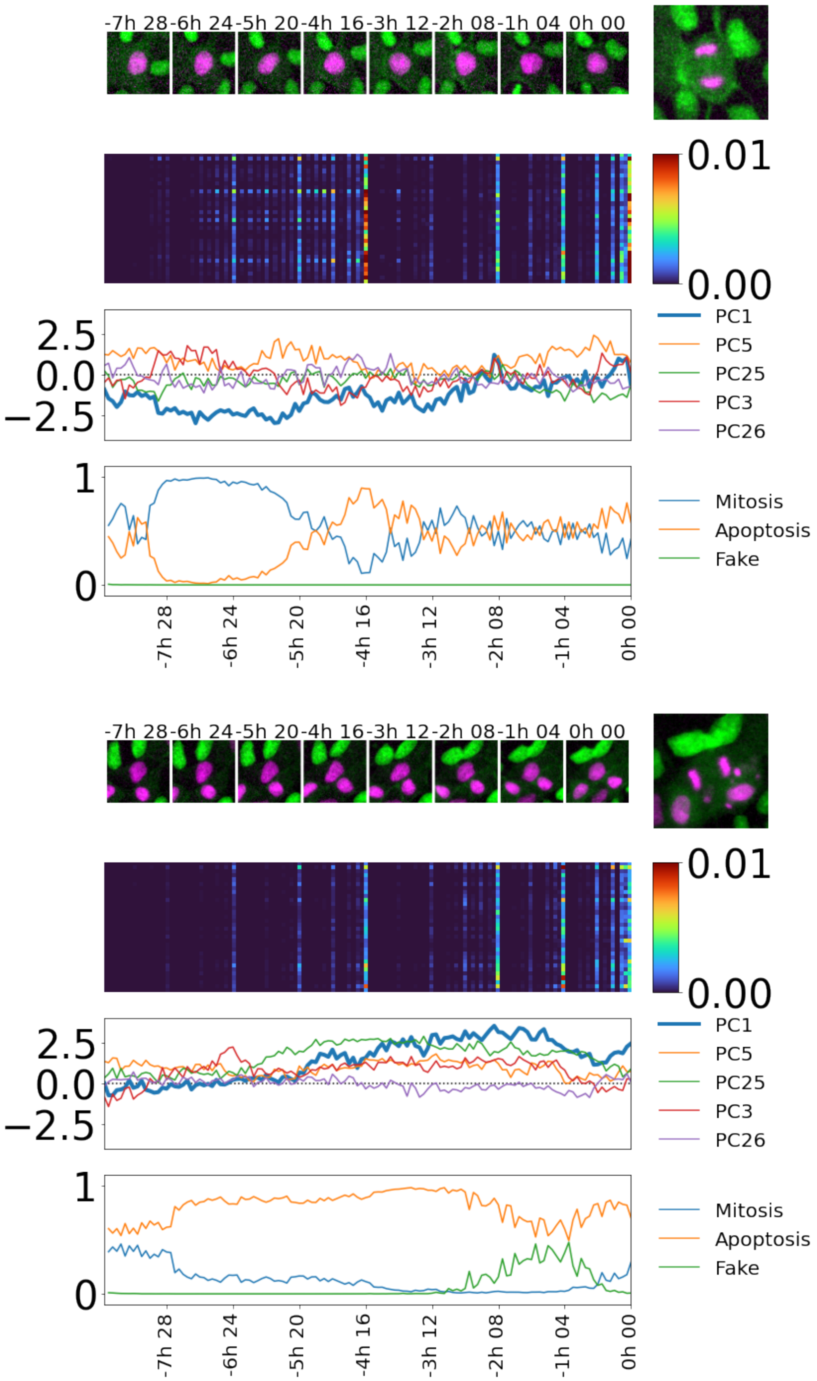

Figure S18: **Example incorrect predictions of cell fate in the BIRB796 treated dataset.** Two example trajectories are shown, with one sub-figure for each. At the top of each sub-figure is placed a collage of time-points of the trajectory before the "cutoff" point. To the right is the final time-point of the trajectory (after the cutoff point), revealing the cell fate. Below that is shown, in order: the confidence plot, showing the TCN's predictions over time; a plot of PC1, the most important principal component for fate prediction; a saliency heat-map showing which components were most important to the prediction, and when and; a saliency plot over time for PC1.

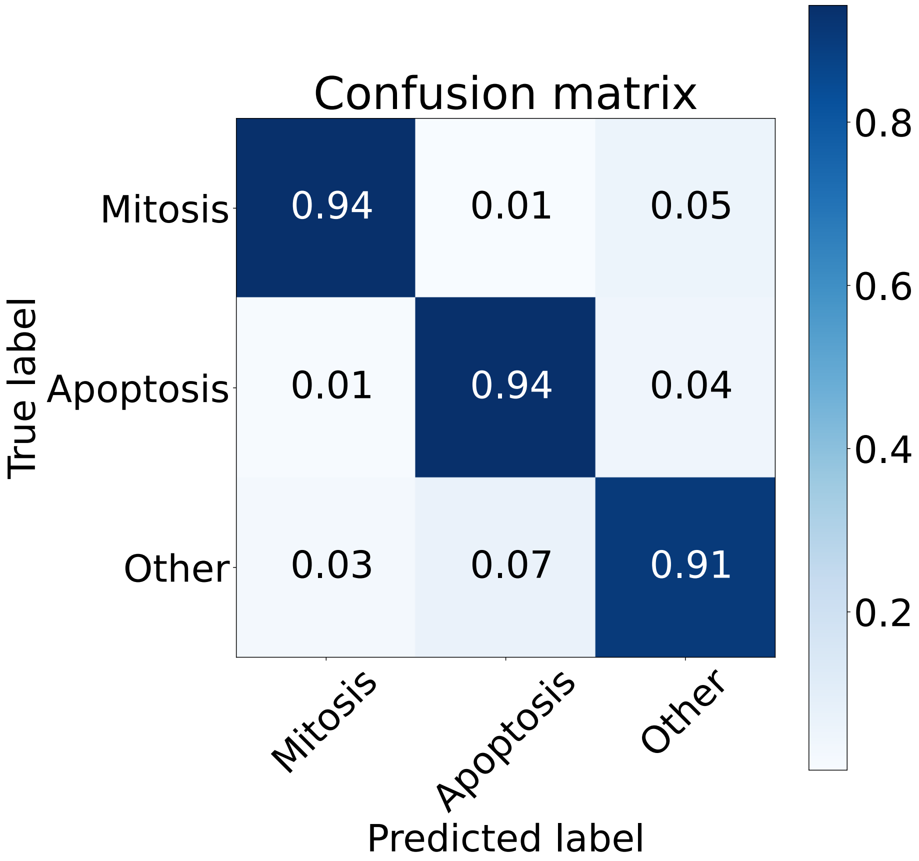

Figure S19: **Normalized confusion matrix for the testing of a neural network trained to discriminate between the final-frame encodings of mitotic, apoptotic and "other" trajectories.** This model was trained for 100 epochs, and its architecture consisted of three dense layers with 64, 128 and 256 units respectively, followed by a final logits layer with 3 units.

### 3 Supplementary tables

Table 1: Datasets used in this study. All movies were captured with a size 1600×1200 pixels (corresponding to 530×400 µm) with 3 channels (Brightfield, GFP and RFP).

| Drug treatment | Cell types | | | Movies | Images |
|---|---|---|---|---|---|
| | GFP | RFP | Seeding Ratio | | |
| None | $MDCK^{WT}$ | $scrib^{kd}$ | 90:10 | 23 | 23,521 |
| | $MDCK^{WT}$ | $scrib^{kd}$ | 50:50 | 62 | 66,607 |
| | $MDCK^{WT}$ | $scrib^{kd}$ | 10:90 | 26 | 26,386 |
| Uninduced | $MDCK^{WT}$ | $scrib^{kd,\ tet-}$ | 50:50 | 6 | 5,925 |
| | $MDCK^{WT}$ | $scrib^{kd,\ tet-}$ | 10:90 | 5 | 5,814 |
| BIRB796 | $MDCK^{WT}$ | $scrib^{kd}$ | 50:50 | 5 | 5,108 |
| Total | | | | 127 | 133,361 |

Table 2: Single-cell trajectory training datasets used in this study. († see Methods)

| Fate | $MDCK^{WT}$ | $scrib^{kd}$ | Total |
|---|---|---|---|
| Apoptotic | 384 | 1841 | 2,225 |
| Mitotic | 34813 | 1249 | 36,062 |
| Fake | Dynamically generated† | | |
| Total | 35197 | 3090 | 38287 |

Table 3: Drugs used in this study

| Name | Biochemical activity | Concentration (µM) | Reference |
|---|---|---|---|
| BIRB796 | p38 kinase inhibitor | 2 | [8] |
| DMSO | control | - | |

Table 4: The testing macro-F1 of trained TCNs using various VAE models.

| Model | $MDCK^{WT}$ | $scrib^{kd}$ | Average | Rank |
|---|---|---|---|---|
| Small-View, All Cells | 91 ± 3% | 84 ± 3% | 87 ± 2% | 1 |
| Mid-View, Central Cell Only | 90 ± 3% | 85 ± 4% | 87 ± 2% | 2 |
| Mid-View, All Cells | 81 ± 3% | 75 ± 4% | 78 ± 2% | 3 |
| Mid-View, Neighbour Cells Only | 78 ± 3% | 74 ± 4% | 76 ± 3% | 4 |
| Large-View, All Cells | 76 ± 3% | 65 ± 5% | 70 ± 3% | 5 |

### 4 Supplementary movies

#### 4.1 Movie S1

Timelapse acquistion and tracking of single cell, showing three different spatial scales extracted to form the glimpse.

#### 4.2 Movie S2

Glimpse extracted from movie S1.

#### 4.3 Movie S3

Example cell detection and tracking for MDCK$^{WT}$:scrib$^{kd}$ dataset.

#### 4.4 Movie S4

Example cell detection and tracking for MDCK$^{WT}$:scrib$^{kd, tet-}$ dataset.

#### 4.5 Movie S5

Example cell detection and tracking for MDCK$^{WT}$:scrib$^{kd}$ + 2 μM BIRB796 dataset.

#### 4.6 Movie S6

Example $\tau$-VAE output for MDCK$^{WT}$:scrib$^{kd}$ dataset.

#### 4.7 Movie S7

Example $\tau$-VAE output for MDCK$^{WT}$:scrib$^{kd, tet-}$ dataset.

#### 4.8 Movie S8

Example $\tau$-VAE output for MDCK$^{WT}$:scrib$^{kd}$ + 2 μM BIRB796 dataset.
